# Pyruvate metabolism dictates fibroblast sensitivity to GLS1 inhibition during fibrogenesis

**DOI:** 10.1101/2024.01.30.577965

**Authors:** Greg Contento, Jo-Anne A Wilson, Brintha Selvarajah, Manuela Platé, Delphine Guillotin, Valle Morales, Marcello Trevisani, Vanessa Pitozzi, Katiuscia Bianchi, Rachel C Chambers

## Abstract

Fibrosis is a chronic disease characterized by excessive extracellular matrix (ECM) production which leads to destruction of normal tissue architecture and disruption of organ function. Fibroblasts are key effector cells of this process and respond to a host of pro-fibrotic stimuli, including notably the pleiotropic cytokine, TGF-β_1_, which promotes fibroblast to myofibroblast differentiation. This is accompanied by the simultaneous rewiring of metabolic networks to meet the biosynthetic and bioenergetic needs of contractile and ECM-synthesizing cells, but the exact mechanisms involved remain poorly understood. In this study, we report that extracellular nutrient availability profoundly influences the TGF-β_1_ transcriptome of primary human lung fibroblasts (pHLFs) and the “biosynthesis of amino acids” emerges as a top enriched transcriptional module influenced by TGF-β_1_. We subsequently uncover a key role for pyruvate in influencing the pharmacological impact of glutaminase (GLS1) inhibition during TGF-β_1_-induced fibrogenesis. In pyruvate replete conditions which mimic the physiological concentration of pyruvate in human blood, GLS1 inhibition is ineffective in blocking TGF-β_1_-induced fibrogenesis, as pyruvate is able to be used as the substrate for glutamate and alanine production via glutamate dehydrogenase (GDH) and glutamic-pyruvic transaminase 2 (GPT2), respectively. We further show that dual targeting of either GPT2 or GDH in combination with GLS1-inhibition is required to fully block TGF-β_1_-induced collagen synthesis. These findings embolden a therapeutic strategy aimed at additional targeting of mitochondrial pyruvate metabolism in the presence of a glutaminolysis inhibitor in order to interfere with the pathological deposition of collagen in the setting of pulmonary fibrosis and potentially other fibrotic conditions.

## 1. Introduction

Fibrosis is the concluding pathological outcome and major cause of morbidity and mortality in a number of common chronic inflammatory, immune-mediated and metabolic diseases [1]. The progressive and relentless deposition of a collagen-rich extracellular matrix (ECM) is the cornerstone of the fibrotic response and often culminates in organ failure and premature death. Despite intense research efforts, there remains a paucity of effective treatment options. Idiopathic pulmonary fibrosis (IPF) represents the most rapidly progressive and lethal of all fibrotic diseases and is associated with a dismal median survival of 3.5 years from diagnosis [2, 3]. Although the approval of pirfenidone and nintedanib for the treatment of IPF signalled a watershed moment for the development of anti-fibrotic therapeutics, these agents slow but do not halt disease progression [4, 5]. There therefore remains a pressing need to identify novel anti-fibrotic therapeutic strategies.

Activated fibroblasts and myofibroblasts are the key effector cells of the fibrogenic response and are responsible for the synthesis and deposition of collagen and other extracellular matrix (ECM) proteins during normal tissue repair, as well as in pathological tissue fibrosis [6]. These cells can originate from several sources, including recruited and resident fibroblasts, which differentiate in response to pro-fibrotic stimuli released at sites of acute and chronic injury, primarily the potent pro-fibrotic cytokine, transforming growth factor β_1_ (TGF-β_1_) [7]. Myofibroblasts are also expanded as a result of the stromal reaction in cancer and current evidence provides strong support that the density of stromal myofibroblasts is associated with poor survival in solid cancers [8].

Metabolic reprogramming is a hallmark of cancer and supports the requirements of exponential growth and proliferation, as well as metastasis as the cancer progresses [9]. This has led to the development of several strategies aimed at exploiting the metabolic vulnerabilities of cancer cells as a potential therapeutic strategy with several metabolism-targeting therapeutic agents demonstrating good tolerability and efficacy in recent clinical trials [10]. One of these is CB-839, a nanomolar-range glutaminase 1 (GLS1) inhibitor that is being investigated in several trials (NCT02071862, NCT02071927, NCT02071888, among others). GLS1 is the rate-limiting enzyme for glutaminolysis, a process which converts glutamine into the anaplerotic intermediate α-ketoglutarate. This process is important for the generation of glutamate which is a major amino group donor for aminotransferase reactions, producing amino acids such as alanine or aspartate.

Metabolic reprogramming is also increasingly recognised as an important feature of pathological fibrosis [11, 12]. Indeed, there is accumulating *in vitro* evidence that fibroblast differentiation in response to TGF-β_1_ is accompanied by the reconfiguration of fibroblast metabolic networks to meet the biosynthetic and energetic needs of cell differentiation and ECM production [13]. Identifying and targeting the synthetic vulnerabilities of myofibroblasts may therefore similarly herald a new era for the development of future anti-fibrotic therapeutic approaches.

Glucose and glutamine metabolism are highly modulated following TGF-β_1_ stimulation yet the concentrations of these two major carbon and nitrogen sources are highly variable among tissue culture media which may not only influence intracellular metabolic networks but also perturb the balance between gene expression and metabolism [14, 15]. Several studies in multiple disease models have shown that altering the concentrations of these key nutrients can be therapeutically beneficial [16, 17]. This strategy is particularly promising in disease contexts where the immediate nutrient supply is unknown or variable throughout pathogenesis, such as a poorly vascularised but growing tumour or fibroblastic foci, the hallmark lesion associated with IPF.

In order to identify novel anti-fibrotic strategies based on the exploitation of key metabolic vulnerabilities of myofibroblasts, we investigated the influence of nutrient availability on the TGF-β_1_-induced transcriptome and collagen synthetic capability of primary human lung fibroblasts. We found that the TGF-β_1_-regulated transcriptome is highly sensitive to nutrient availability and further discover a novel role for pyruvate in terms of dictating the sensitivity of fibroblasts to GLS1 inhibition during fibrogenesis. This effect was traced to a superior ability for exogenous pyruvate over endogenously produced pyruvate from glucose to maintain amino acid biosynthesis through the enzymes glutamic pyruvic transaminase 2 (GPT2) and glutamate dehydrogenase (GDH). The findings reported here build a case for the potential of anti-fibrotic approaches involving the inhibition of GLS1 in combination with inhibition of either GPT2 or GDH to target key metabolic vulnerabilities of hyper-synthetic activated fibroblasts during fibrogenesis.

## 2. Materials and methods

### 2.1 Cytokines, metabolites and compounds

Only cytokine used was TGF-β_1_ (R&D Biosystems). Pharmacological inhibitors used were CB-839 (MedChemExpress), EGCG (Sigma) and L-Cycloserine (Sigma). For metabolites, dimethyl 2-oxoglutarate, L-glutamic acid dimethyl ester hydrochloride, L-alanine, L-proline. sodium pyruvate, sodium lactate, and D-(+)-glucose were purchased from Sigma and L-glutamine from ThermoFisher.

### 2.2 Primary human lung fibroblast isolation and culture

Primary human lung fibroblasts (pHLFs) were produced through explant culture and tissue obtained with informed and signed consent and with institutional research ethics committee approval from the UCL Research Ethics Committee (12/EM/0058). Cells were grown in standard DMEM free from glucose and glutamine which were added individually to meet physiological concentrations of 5mM and 0.7mM, respectively or in standard DMEM with 25mM glucose, 1mM pyruvate and 2mM glutamine. Media was supplemented with 10% fetal bovine serum (FBS) (Sigma-Aldrich) and 1% Penicillin (100U/ml)/Streptomycin (100 µg/ml) (Thermo-Fisher). All cultures were routinely screened for mycoplasma.

### 2.3 Transcriptomic analysis by RNA sequencing (RNA-Seq)

Fibroblasts were grown to confluency in 6-well plates and total RNA extracted using RNeasy Kit (Qiagen). Poly(A)-tailed RNA enrichment and library construction was performed using KAPA stranded mRNA-Seq Kit with KAPA mRNA capture beads (KAPABiosystems). High-throughput sequencing was performed at UCL Genomics using the NextSeq sequencing platform (Illumina) and the sequencing data were uploaded to the Galaxy web platform for analysis. Sequenced reads were quality tested using FASTQC (version 0.72) and aligned to the hg38 human genome using HISAT2 with default parameters (version 2.1.0). FeatureCounts was used to quantify counts over reference genes (version 1.6.4) and differential gene expression was carried out using DESeq2 (version 2.11.40.6). Differentially expressed genes were defined as having a p-adjusted value ≤ 0.05 and a log_2_ fold change of ≤-0.58 or ≥0.58 when comparing two experimental conditions. Pathway enrichment analysis was carried out on normalized counts using clusterProfiler (Version 3.17) and GSEA (version 4.1.0) with 1000 permutations, a gene set size filter of 15-500 and the default metric for ranking genes.

### 2.4 Quantification of amino acids using HPLC

Free amino acids were quantified using high-performance liquid chromatography (HPLC). Fibroblast monolayers were washed once with ice-cold PBS before addition of ice-cold RIPA buffer (ThermoFisher Scientific) supplemented with protease inhibitors (cOmpleteMini, Roche). Lysates were then transferred to 1.5ml Eppendorf tubes and vortexed vigorously for 30s prior to centrifugation at 21,000g for 10 minutes at 2°C. Supernatants were then transferred to 3 kDa protein-filter columns (Sigma-Aldrich) and centrifuged at 12,000g for 45 minutes. The flow-through was then dried in a speed-vac and derivatized with 7-chloro-4-nitrobenzo-2-oxa-1,3-diazole (Acros Organics, ThermoFisher Scientific) prior to reverse-phase HPLC (Agilent 1100 series, Agilent Technologies) for hydroxproline isolation using acetonitrile as the organic solvent in a LiChrospher, 100 RP-18 column.

### 2.5 Quantification of hydroxyproline from cellular supernatants using HPLC

Fibroblast pro-collagen secretion was assessed by high-performance liquid chromatography (HPLC) quantification of hydroxyproline in cell supernatants. Proteins were precipitated by adding ethanol to a concentration of 66% (v/v), vortexed and left overnight at 4°C. The solutions were then filtered using 0.45 µm filters (Millipore) which were then incubated with 6M hydrochloric acid at 110°C for 16 hours. Samples were then dried at 100°C and derivatized with 7-chloro-4-nitrobenzo-2-oxa-1,3-diazole (Acros Organics, ThermoFisher Scientific) prior to reverse-phase HPLC (Agilent 1100 series, Agilent Technologies) for hydroxproline isolation using acetonitrile as the organic solvent in a LiChrospher, 100 RP-18 column.

### 2.6 High-Content Imaging for collagen type I deposition analysis

Collagen type I deposition was measured in 96-well plates using high-content immunofluorescence-based imaging in molecular crowding conditions as previously described [18]. Briefly, pHLFs were grown in a 96-well plate in 10% FBS media until confluency before replacement with 0.4% FBS media for 24 hours. Media was then replaced once more with 0.4% FCS containing ascorbic acid (16.6µg/ml) and Ficoll 70 and Ficoll 400. Cells were then treated with compounds for 2 hours prior to TGF-β_1_ stimulation (1ng/ml, R&D systems. After 48 hours, the media was removed and monolayers fixed with methanol prior to antibody addition targeting human collagen I (Sigma) followed by AlexaFluor 488 secondary antibody (Life Technologies) and nuclear DAPI counterstain. Fluorescence was quantified using an ImageXpress Micro XLS high-content imaging system at 20x magnification (Molecular Devices) with 4 fields of view per well and collagen signal normalised to cell count.

### 2.7 Immunoblotting

pHLFs were lysed using ice-cold PhosphoSafe buffer (Merck Millipore) supplemented with protease inhibitors (cOmpleteMini, Roche). Protein concentration was determined using a Pierce BCA protein assay kit (ThermoFisher Scientific) and 10 µg protein loaded into 15-well NuPage 4-12% Bis-Tris gels (ThermoFisher Scientific) for SDS PAGE separation. Gels were then transferred to nitrocellulose and protein levels assessed by Western blotting with the following antibodies: GLS1 (12855-1-AP), GPT2 (HPA051514), GOT2 (PA5-27572), GOT1 (CST#34423), P5CS (ab223713), SHMT2(CST#12762), ASNS (14681-1-AP), GDH (14299-1-AP), PSAT1 (PA522124) and α-tubulin (CST#9099). Dilutions were 1:1000 except for α-tubulin which was 1:5000. All densitometries are presented relative to α-tubulin unless stated otherwise.

### 2.8 siRNA-mediated protein expression knock-down

pHLFs were seeded in a 12-well plate in 10% FBS media for 24-hours before transfection with 25nM siRNAs (Dharmacon, SMARTpool) targeting *GPT2*, *GOT1*, *GOT2, PSAT1, SHMT2, ASNS, GDH* and *ALDH18A1* using RNAiMax lipofectamine (Invitrogen) following the manufacturer’s recommendations. The following day the media was replaced with 0.4% FBS media and left for a further 24-hours before TGF-β_1_ treatment.

### 2.9 Metabolomic analysis and labelling using LC-MS

pHLFs were plated in 6-well plates and grown to confluency. For labelling studies, ^13^C_6_-Glucose or ^13^C_3_-pyruvate were added for either the full 48 hours or in the last 8 hours accompanied by a media change (Cambridge Isotope Laboratories). Cells were then washed with ice-cold PBS three times and metabolites extracted using an extraction solvent consisting of 50% methanol, 30% acetonitrile and 20% ultrapure water added with 100 ng/ml of HEPES. Cells were scraped and transferred to a shaker for 15 minutes at 4°C before an incubation at −20°C for one hour. Samples were then spun at maximum speed for 10 minutes at 4°C and supernatants transferred to new tubes (process repeated twice). Supernatants were then transferred to autosampler glass vials and stored at −80°C until analysis. Samples were then loaded onto the machine in a randomised order to avoid machine drift bias. LC-MS analysis was performed using a Q Exactive Hybrid Quadrupole-Orbitrap mass spectrometer coupled to a Dionex U3000 UHPLC system (Thermo Fisher Scientific). A Sequant ZIC-pHILIC column (150 mm x 2.1 mm) was used with a guard column (20 mm x 2.1 mm) (Merck Milipore). The temperature was maintained at 45°C and mobile phase consisted of 20 mM ammonium carbonate and 0.1% acetonitrile. Flow rate was 200 µl/min with previously described gradient [19]. The mass spectrometer was used in full MS and polarity switching mode. Spectra were analysed using Xcalibur Qual Browser and Xcalibur Quan Browser software (Thermo Fisher Scientific). The mzCloud advanced mass spectral database for used for the untargeted analysis.

### 2.10 mRNA quantification using real-time quantitative PCR (qPCR)

RNA was extracted using RNeasy Mini kit (Qiagen) following the manufacturer’s instructions. Real-time PCR was performed using a Mastercycler Realplex ep gradient S (Eppendorf). Amplification was performed for 40 cycles with a melting temperature of 95C for 15 seconds and an annealing temperature of 60C for one minute. Relative quantification was derived using 2-ΔCt (ΔCt calculated from the mean of two normalising genes, *ATP5B* and *B2M* (PrimerDesign Ltd, primer sequence proprietary). The following primers were used: GPT2 forward 5’-TCCTCACGCTGGAGTCCATGA-3’ and reverse 5’-ATGTTGGCTCGGATGACCTCTG-3’; GDH (GLUD1) forward 5’-CTCCAAACCCTGGTGTCATT-3’ and reverse 5’-CACACGCCTGTGTTACTGGT-3’.

### 2.11 Statistics

All data are expressed as the means +/-SD and generated using GraphPad Prism version 8.0 and all experiments repeated at least 3 times. Statistical differences between two groups was determined using one-way or 2-way ANOVA with Tukey post-hoc test. Four-parameter and non-linear regression analysis was used to produce IC_50_ values from compound concentration curves. The α level was 0.05 for all tests.

### 2.12 Study approval

Primary human lung fibroblasts (pHLFs) were produced through explant culture and tissue obtained with informed and signed consent and with institutional research ethics committee approval from the UCL Research Ethics Committee (12/EM/0058).

### 2.13 Data availability

The RNA-Seq dataset has been deposited and will be publicly available at GSE244098. All other data needed to evaluate the conclusions in the paper are present in the paper, the supplementary data or publicly available datasets in gene expression omnibus (GEO) as specified.

## 3 Results

### 3.1 The TGF-β_1_-induced transcriptome is modulated by glucose and glutamine metabolism

TGF-β_1_ promotes a widespread transcriptional response in multiple cell types and has been shown to increase chromatin accessibility of genomic enhancer regions by as much as 80% [20]. Chromatin accessibility is intimately dependent on metabolic reactions, including epigenetic acetylation, methylation and demethylation which are all highly dependent on glucose and glutamine metabolism. However, the concentrations of glucose (Glc) and glutamine (Gln) vary widely in the culture media employed across mechanistic studies aimed towards understanding these processes in health and disease. DMEM is a common media formulation for the cultivation of fibroblasts and traditionally contains 25 mM Glc and 2 mM Gln (termed DMEM^High^ onwards), far higher than the approximate 5 mM Glc and 0.7 mM Gln (termed DMEM^Low^ onwards) present in DMEM formulations aimed at replicating human sera levels. Pyruvate concentrations also differ from absent (DMEM^Low^) to 1mM (DMEM^High^). To investigate the effects of media composition on the TGF-β_1_ transcriptome, we compared an RNA-Seq dataset performed in primary human lung fibroblasts (pHLFs) cultured in DMEM^Low^ with a previously published dataset of the same cells exposed to the same concentration of TGF-β_1_ (1ng/ml) in DMEM^High^ from our laboratory (GSE102674).

Differential gene expression analysis revealed substantial differences between the total number of significantly modulated genes and the magnitude of the fold changes induced by TGF-β_1_, with DMEM^High^ leading to nearly 3-fold more differentially expressed genes (DEGs) than DMEM^Low^ (Fig. 1a-c). The high number of unique DEGs in DMEM^High^ (2,986) led us to examine metabolic pathway expression to determine if there was any transcriptional evidence of the influence of high carbon and nitrogen sources available in this media. KEGG pathway analysis indeed showed central carbon metabolism as a top hit (p=0.0024) which included genes involved in glycolysis, such as lactate dehydrogenase A (*LDHA*) and pyruvate dehydrogenase E1 subunit beta (*PDHB*) (Supplementary Fig. 1). We next wanted to ensure that regardless of the media, the central processes involved in TGF-β_1_ signalling (i.e. ECM remodelling) were retained and significantly modulated. Reactome analysis on these 1,279 shared genes was used to determine these processes and this revealed terms such as collagen biosynthesis, fibril formation and assembly, ECM formation and tRNA aminoacylation (the latter potentially to meet the enhanced protein synthetic demands of TGF-β_1_-activated hyper-synthetic fibroblasts) (Fig. 1d). The ability of both media formulations to support a robust TGF-β_1_ procollagen response at the protein level was confirmed by assessing hydroxyproline levels in cell supernatants, with DMEM^High^ cultured cells showing a slightly higher response (Fig. 1e).

**Figure 1.**
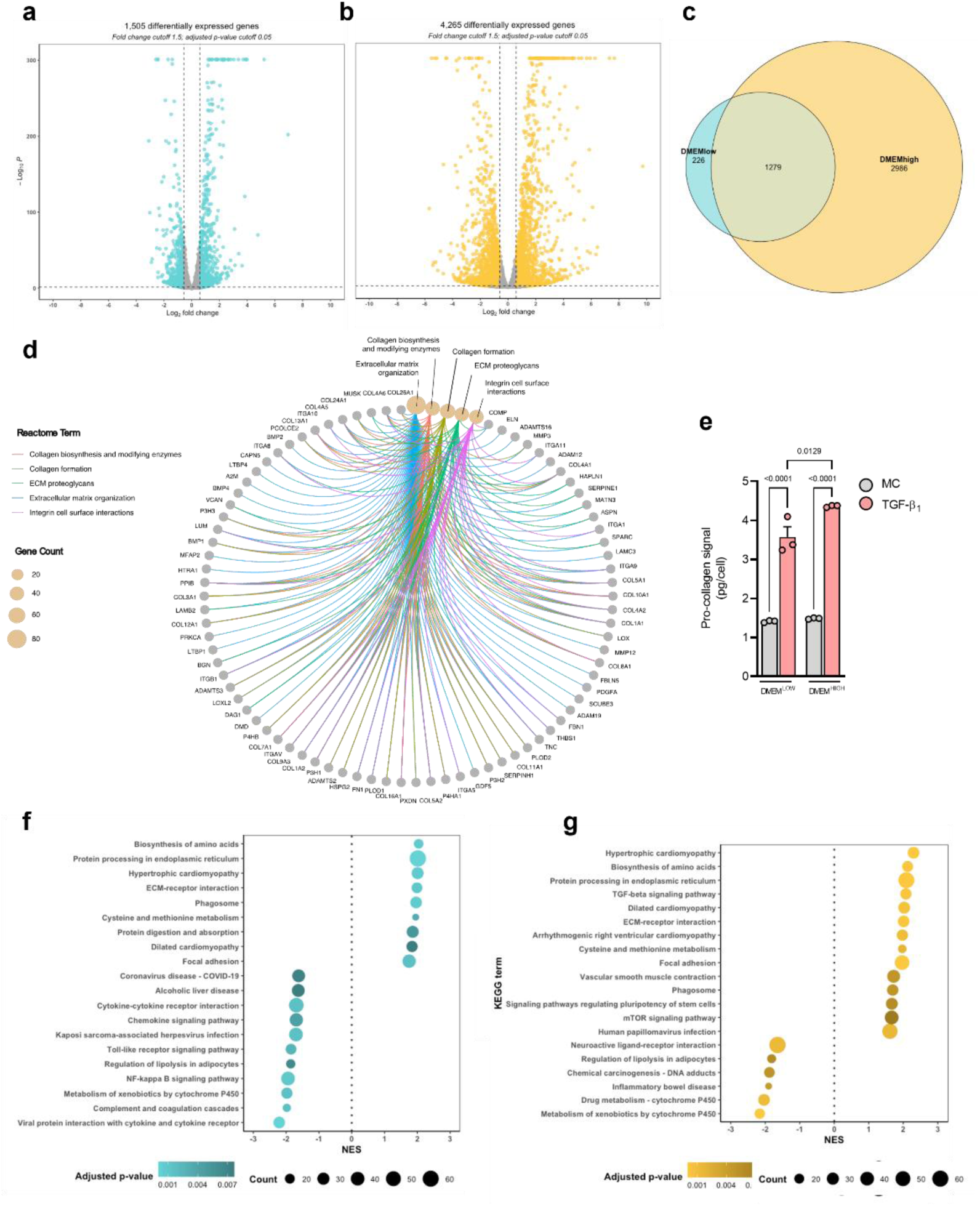
The TGF-β_1_induced transcriptome is sensitive to media composition. **a,b** Volcano plots showing differentially expressed genes in a DMEM^Low^ and b DMEM^High^ of pHLFs stimulated with TGF-β_1_ (1 ng/ml) for 24 h with cut-off values q < 0.05 and fold-change ± 1.5. **c** Overlap diagram of DEGs between DMEM^Low^ and DMEM^High^. **d** Network plot of enriched reactome terms of DEGs shared between DMEM^Low^ and DMEM^High^. **e** Supernatants collected from pHLFs stimulated with TGF-β_1_ (1 ng/ml) for 48 h and hydroxyproline quantified using HPLC (*n*=3) and is representative of 5 independent experiments, f, g Dot-plots showing top 20 enriched KEGG terms in **f** DMEM^Low^and **g** DMEM^High^. Data are presented as mean ± SD and differences evaluated between groups with two-way ANOVA with tukey multiple comparison testing.

We next performed gene-set enrichment analysis (GSEA) in both media formulations shown in figures 1a and 1b. This analysis revealed that almost half of the top 20 gene sets were unique to the media used (Fig. 1f-g). Interestingly, this analysis identified the ‘biosynthesis of amino acids’ as a top TGF-β_1_-dependent module in both media conditions, indicating that even with markedly higher nutrients sources present in DMEM^High^, this pathway was highly modulated by TGF-β_1_ in both nutrient environments.

### 3.2 TGF-β_1_-modulated intracellular amino acid pools influence metabolic vulnerabilities

Amino acid biosynthesis is known to be highly dependent on the availability of glutamine which provides transferrable amino groups to α-ketoacids after its conversion to glutamate (Fig. 2a). Given that the glutamine concentration is approximately 3-fold higher in DMEM^High^ (2mM versus 0.7mM in DMEM^Low^), we measured the intracellular levels of glutamate-dependent amino acids following TGF-β_1_ stimulation. We found that TGF-β_1_ decreased intracellular Gln levels in DMEM^Low^ by more than 90%, to levels barely above the limit of detection by 48 hours; this was in stark contrast to the high levels of intracellular Gln measured in DMEM^High^ at baseline and in response to TGF-β_1_ (Fig. 2b). We questioned whether this marked decrease in intracellular Gln levels in TGF-β_1_-treated cells in DMEM^Low^ was due to a depletion of extracellular Gln levels over the 48-hour culture period. However, we found that extracellular Gln levels had only dropped by close to 50% (377 µM) at this time-point (Supplementary Fig. 2a). TGF-β_1_ increased the intracellular levels of all other amino acids measured and this effect was more pronounced in DMEM^High^ (except for aspartate levels, which were unchanged by TGF-β_1_). Since glutamate, asparagine, aspartate, alanine and proline are not added to these DMEM formulations, the increased intracellular levels suggest increased amino acid biosynthesis induced in response to TGF-β_1_ rather than increased uptake from the media. In support of this, we found that TGF-β_1_-induced levels of serine and glycine, which are both present in DMEM, were similar for DMEM^Low^ and DMEM^High^. Intracellular levels of glutamate in unstimulated cells cultured in DMEM^High^ were higher than the glutamate levels in TGF-β_1_-treated cells cultured in DMEM^Low^. This might potentially indicate an enhanced capacity of fibroblasts in DMEM^High^ to synthesize non-essential amino acids owing to a larger pool of available intracellular glutamate.

**Figure 2.**
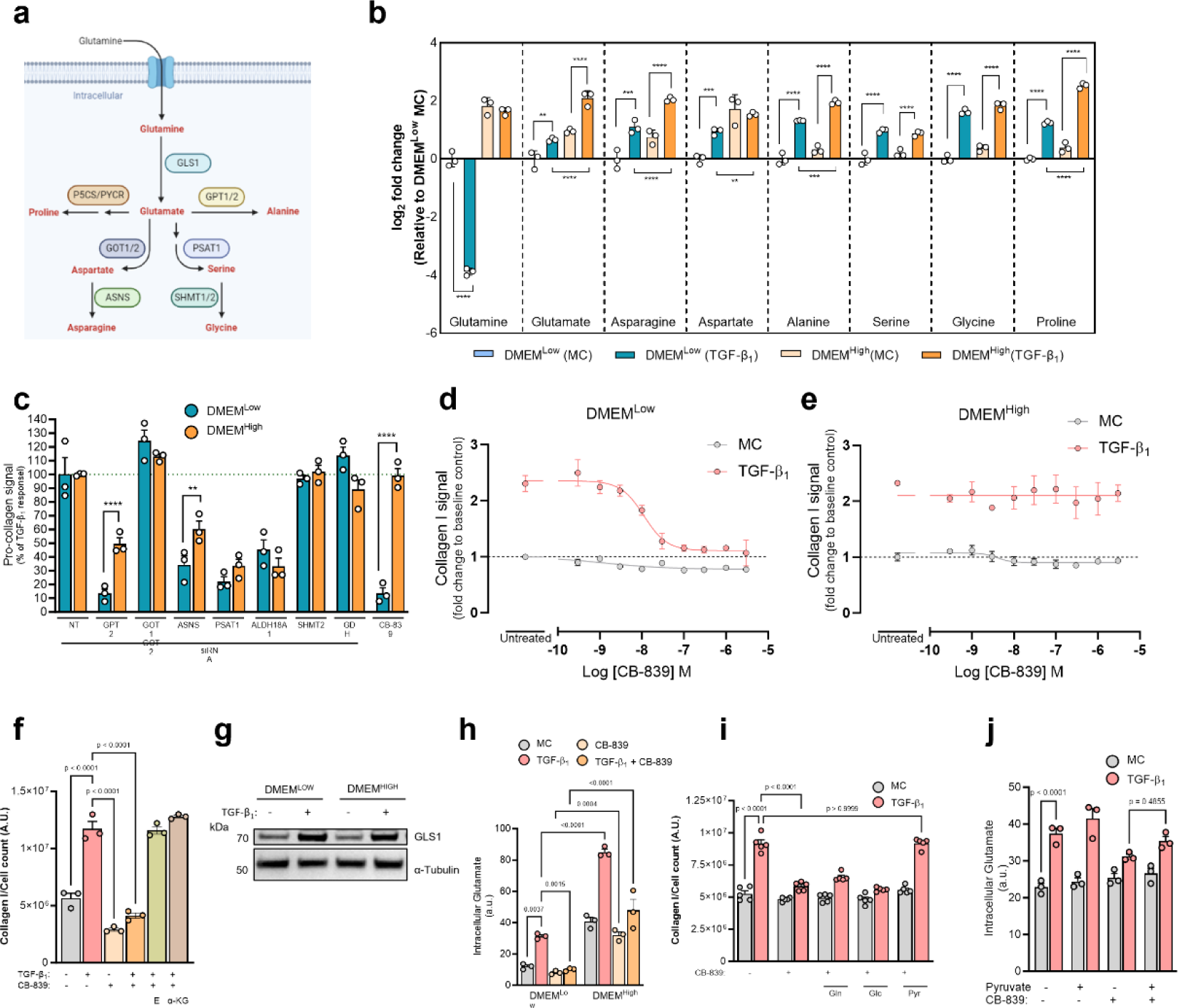
TGF-β_1_-modulated intracellular amino acid pools influence metabolic vulnerabilities. **a** Schematic showing amino acid biosynthetic pathways which utilise glutamine-derived glutamate, **b** Intracellular levels of amino acids in pHLFs in DMEM^Low^ or DMEM^High^ stimulated with TGF-β_1_ (1 ng/ml) for 48 h quantified using HPLC (*n*=3). **c** Supernatants collected from pHLFs stimulated with TGF-β_1_(1 ng/ml) for 48 h following transfection with non-targeting siRNA for control and CB-839 treated groups (1 μM of CB-839) or siRNA targeting GPT2, GOT1 and GOT2, ASNS, PSAT1, ALDH18A1, SHMT2 or GDH and hydroxyproline quantified using HPLC (*n*=3). Data is normalized to non-targeting control stimulated with TGF-β_1_, (1 ng/ml). **d,e** pHLFs were grown in DMEM^Low^ or DMEM^High^ and pre-incubated with increasing concentrations of CB-839 for 1 h before being stimulated with TGF-β_1_, (1 ng/ml) for 48 h and collagen I deposition assessed by macromolecular crowding assay. Data are expressed as collagen I signal as a fold-change of the media control (0.1% DMSO) (*n*=4 fields of view imaged per well) normalised to cell count which was obtained by staining nuclei with DAPI. **f** Collagen deposition quantified 48 hours after TGF-β_1_, (1 ng/ml) stimulation from pHLFs growing in DMEM^Low^ and 1 μM CB-839 with supplementation of dimethylated forms of glutamate (E, 4mM) and a-KG (4mM). **g** Immunoblot of intracellular protein lysates from pHLFs 24 hours following TGF-β, stimulation (1 ng/ml) grown in DMEM^Low^ or DMEM^High^. **h** pHLFs were grown in DMEM^Low^ or DMEM^High^ and stimulated with TGF-β_1_, (1 ng/ml) and CB-839 (1μM) for 48 h before being lysed and glutamate measured by HPLC (*n*=3). **i** pHLFs were grown in DMEM^Low^supplemented with glutamine (1.3 mM), glucose (20 mM) or pyruvate (1 mM) before pre-incubation with 1 pM CB-839 for 1 h before TGF-β_1_, (1 ng/ml) stimulation for 48 h and collagen I deposition assayed by macromolecular crowding assay. **j** Intracellular glutamate levels in pHLFs grown in DMEM^Low^ supplemented with pyruvate (1 mM) and pre-incubation with 1 pM CB-839 for 1 h before TGF-β_1_, (1 ng/ml) stimulation for 48 h and quantification achieved using HPLC (*n*=3) Data are presented as mean + SD and differences evaluated between groups with two-way ANOVA with tukey multiple comparison testing.

We next investigated whether the enzymes involved in amino acid biosynthesis were conditionally essential dependent on media composition for TGF-β_1_-induced collagen synthesis. At the time of this study GLS1 was the only enzyme with a pharmacological inhibitor with high selectivity and nanomolar potency (CB-839) [21]. We used this compound to address the role of GLS1 and we employed siRNA-mediated gene knockdown for all other enzymes. We opted to silence both the cytoplasmic and mitochondrial forms of GOT, *GOT1* and *GOT2*, respectively, since both can generate aspartate and may therefore compensate for one another (Supplementary Fig. 2b). These studies revealed GPT2, ASNS and GLS1 as being more critical enzymes to enable fibroblasts in DMEM^Low^ than in DMEM^High^ to mount a TGF-β_1_ collagen response (Fig. 2c). We further found that PSAT1 and ALDH18A1 (gene product is P5CS) were crucial for TGF-β_1_-induced collagen levels regardless of DMEM formulation, which is in line with previous work highlighting the important role of serine and proline biosynthetic pathways in TGF-β_1_-induced collagen synthesis [22, 23]. We defined the mechanism of action of GPT2, ASNS and ALDH18A1 to the absence in DMEM of alanine, asparagine and proline, respectively, as introduction to the growth media of each respective amino acid completely rescued TGF-β_1_-induced collagen levels in DMEM^Low^ (Supplementary Fig. 2c,d). It is therefore plausible that the greater stores of intracellular alanine and asparagine of fibroblasts in DMEM^High^ may be attributable for the differences in effect of collagen production observed for siGPT2 and siASNS, respectively, when compared to DMEM^Low^.

Unlike siGPT2, siASNS or siALDH18A1 treatment which reduced TGF-β_1_ collagen levels in both DMEM formulations, inhibition of GLS1 completely prevented TGF-β_1_ collagen synthesis in DMEM^Low^ but failed to produce any effect in DMEM^High^ (Fig. 2d,e). GLS1 is the rate-limiting enzyme for glutaminolysis, which produces glutamate for subsequent α-ketoglutarate generation into the TCA cycle. We examined this mechanism of action by supplementing fibroblasts with a cell-permeable form of either glutamate or α-ketoglutarate where both individually restored the TGF-β_1_-induced collagen response under GLS-1 inhibited conditions (Fig. 2f). The media difference in impact of CB-839 on the TGF-β_1_-induced collagen response was not the result of different GLS1 protein abundance in the respective media formulations, since the GLS1 splice variants, KGA and GAC, were induced to similar levels by TGF-β_1_ treatment in DMEM^High^ and DMEM^Low^ (Fig. 2g). Moreover, intracellular TGF-β_1_-stimulated glutamate levels were GLS1-dependent in both media, but cells cultured in DMEM^High^ retained a much higher abundance of glutamate following CB-839 exposure (Fig. 2h). We therefore explored the possibility that the high levels of Gln present in DMEM^High^ may allow for a higher glutamate pool which is sufficient for cells to mount a full TGF-β_1_ collagen response without GLS1 activity. However, when examining the cause for this difference by individually adjusting the concentrations of the three components which differ between the DMEM formulations (glucose, pyruvate and glutamine) in DMEM^Low^ to the levels present in DMEM^High^, we found that it was pyruvate and not Gln (as expected) which was responsible for restoring the TGF-β_1_-induced collagen response in GLS1-inhibited cells cultured in DMEM^Low^ (Fig. 2i). These observations suggest that pyruvate confers resistance to GLS1 inhibition in DMEM^High^, an effect that was observed from 100 µM (physiological human blood concentration) onwards by addition of pyruvate to DMEM^Low^-cultured cells (Supplementary Fig. 2e). Furthermore, intracellular glutamate levels were not significantly increased by the presence of pyruvate and were below the levels measured in DMEM^High^ (Fig. 2j). These data suggest that while the higher glutamate levels were likely a result of high Gln availability, pyruvate rendered cells GLS1-independent in terms of the TGF-β_1_-induced collagen response by affecting another mechanism(s) rather than by influencing glutamate synthesis directly.

### 3.3 Exogenous pyruvate supports the TCA cycle under GLS1 restriction

Depriving cancer cells of Gln-derived glutamate via the inhibition of GLS1 is being actively pursued as a therapeutic strategy in the oncology setting and has also been previously explored as a novel anti-fibrotic approach in pre-clinical studies [24–26]. Pyruvate is the end-product of glycolysis and is also a source of cytosolic NAD+ through lactate synthesis and a major anaplerotic substrate. The most immediate functional overlap between pyruvate metabolism and GLS1-directed metabolism resides in the tricarboxylic acid (TCA) cycle (Fig 3a). We therefore sought to further understand the gene-nutrient interaction between GLS1 and pyruvate in DMEM^Low^ to potentially uncover mechanistic overlaps which may aid in the future translation of therapeutic approaches involving GLS1 antagonists in the setting of fibrosis. We first explored the fate of endogenously produced pyruvate into lactate and acetyl-coA with fully [^13^C]-labelled Glc under GLS1-restricted conditions using liquid-chromatography-mass spectrometry (LC-MS). TGF-β_1_ decreased intracellular Glc levels but this was likely due to increased glycolytic flux as a result of a higher pyruvate pool which was found to be completely m+3 labelled (Fig. 3b,c). Interestingly, CB-839 treatment prevented this increase in Glc-derived pyruvate. We also observed higher Glc levels in control fibroblasts with CB-839 treatment, yet this did not translate to higher pyruvate synthesis. This suggests an inability for cells to effectively metabolise Glc into pyruvate under GLS1 inhibited conditions. Indeed, with CB-839 treatment we found that even in the presence of 25 mM Glc (the concentration found in DMEM^High^), which increased intracellular Glc levels by more than 13-fold compared with 5 mM Glc present in DMEM^Low^, pyruvate levels were unchanged (Supplementary Fig. 3a,b). This decrease in pyruvate levels by CB-839 translated into a lower m+3 lactate pool yet the labelling percentage was unchanged, indicative of the pyruvate contribution to lactate (Fig. 3d). We investigated if this decrease in lactate could impact on TGF-β_1_-induced collagen levels. We supplemented the media with lactate prior to CB-839 treatment but found no changes in the diminished collagen response (Supplementary Fig. 2c). Additionally, we found that with 25 mM glucose (which did not modulate the effectiveness of CB-839 on TGF-β_1_ collagen levels) the lactate pool was fully restored, further indicating that lactate generation was not a mechanism by which pyruvate was working to rescue TGF-β_1_ collagen production (Fig. 2h) (Supplementary Fig. 3d).

**Figure 3.**
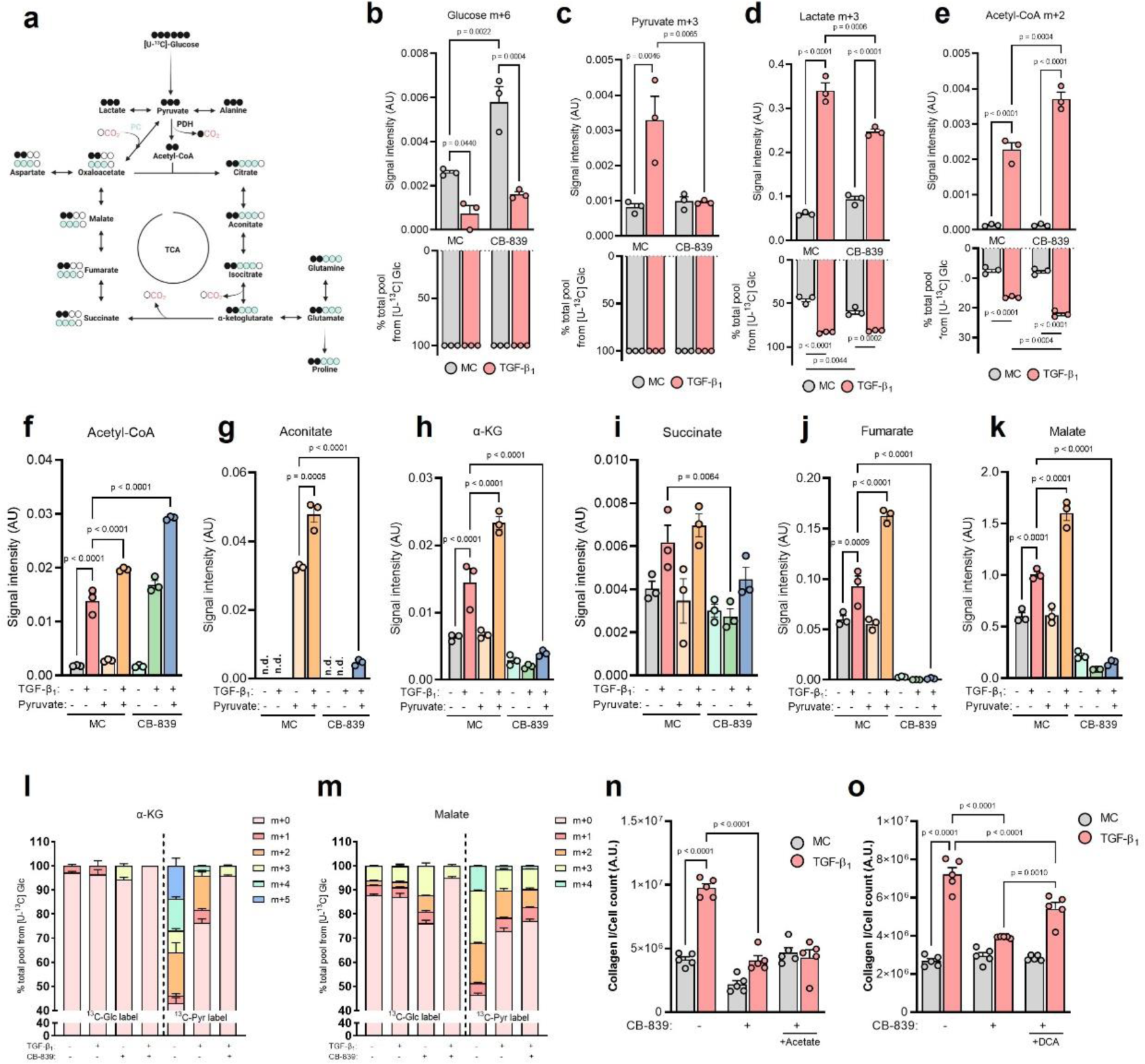
Exogenous pyruvate supports the TCA cycle under GLS1 restriction. **a** Schematic showing U-^13^C-glucose routing to pyruvate and relevant TCA cycle intermediates and amino acids **b-e** Intracellular isotopologue levels and fractional enrichment of specified metabolite in pHLFs grown in DMEM^Low^ supplemented with U-^13^C-glucose (5 mM) and pre-incubated with media control (0.1% DMSO) or 1 μM CB-839 for 1 h before stimulation with TGF-β_1_, (1 ng/ml) for 48 h and quantification achieved using LC-MS (*n*=3). **f-m** Intracellular levels of specified metabolite or **l,m** fractional enrichment in pHLFs grown in DMEM^Low^ supplemented with U-^13^C-glucose (5 mM) or U-^13^C-pyruvate (1 mM) and pre-incubated with media control (0.1% DMSO) or 1 μM CB-839 for 1 h before stimulation with TGF-β_1_ (1 ng/ml) for 48 h and quantification achieved using LC-MS (*n*=3). n,o Collagen deposition quantified 48 hours after TGF-β_1_ (1 ng/ml) stimulation from pHLFs growing in DMEM^1^™ and 1 μM CB-839 with supplementation of acetate (1 mM) or dichloroacetate (DCA, 10 mM).

We next focused on acetyl-coA levels and found that the m+2 signal and the label percentage, indicative of pyruvate dehydrogenase (PDH) activity, was significantly increased by TGF-β_1_ in DMEM^Low^ (Fig. 3e). Inhibition of GLS1 increased these changes, suggesting that cells adapt to the inhibition of glutaminolysis, a major anaplerotic pathway, by enhancing Glc routing into the TCA cycle via acetyl-coA generation. We therefore explored the possibility that exogenous pyruvate may support this compensatory mechanism to levels necessary to sustain a full TGF-β_1_ collagen response. We found that pyruvate supplementation significantly increased TGF-β_1_-induced acetyl-coA levels and increased these levels even further under GLS1 inhibition, indicative of heightened PDH activity (Fig. 3f). Even though we did not measure a decrease in acetyl-coA levels following CB-839 treatment, this result suggested that the overflow of acetyl-coA may be compensating for a potentially decreased contribution from glutaminolysis to the TCA cycle. Indeed, we found the levels of intermediates of the TCA cycle significantly decreased following GLS1 inhibition (Fig. 3g-k). However, supplementation with pyruvate did not alleviate this depression of the TCA cycle. Interestingly, we were only able to detect aconitate in fibroblasts exposed to extracellular pyruvate (Fig. 3g).

To examine whether pyruvate was feeding into the TCA cycle as a carbon substrate we traced the fate of [^13^C]-labelled pyruvate into α-ketoglutarate and malate. We found that exogenous pyruvate contributed more than Glc-derived pyruvate to the total pools of both of these TCA cycle metabolites in both control and TGF-β_1_ conditions (Fig. 3l,m). GLS1 inhibition completely prevented Glc routing into α-ketoglutarate whilst exogenous pyruvate could maintain an m+3 pool. GLS1 inhibition also decreased the m+2 and m+3 fractions of Glc-derived malate, an effect that was not observed with supplemented pyruvate. This suggested that Glc-derived pyruvate had lower flux into the TCA cycle than exogenous pyruvate following GLS1 inhibition and that although the total amounts of TCA cycle intermediates were significantly decreased by CB-839, there existed an active route for pyruvate to sustain both PDH and pyruvate carboxylase (PC) activity. To test this further we provided fibroblasts with acetate as a means of elevating acetyl-coA levels. However, we found this did not alleviate TGF-β_1_-induced collagen levels decreased by CB-839 (Fig. 3n). Given that acetate conversion to acetyl-coA is an ATP dependent process and that the TCA cycle was severely depressed by GLS1 inhibition, we explored whether increasing PDH activity, which uses NAD+, via dichloroacetate (DCA), a potent inhibitor of all pyruvate dehydrogenase kinases, would rescue the TGF-β_1_ collagen response. DCA treatment in CB-839 treated cells significantly increased TGF-β_1_-induced collagen levels to roughly 57% of the TGF-β_1_ response, compared to 22% without DCA treatment (Fig. 3o). These results suggested that at least part of the rescue mechanism(s) of pyruvate on CB-839-inhibited TGF-β_1_-induced collagen levels reside in propping up the TCA cycle and that enhancing pyruvate flux into the TCA cycle can modulate CB-839 sensitivity.

### 3.4 Exogenous pyruvate supports amino acid biosynthesis under GLS1 restriction

The partial rescue of DCA on CB-839-decreased collagen production suggested that exogenous pyruvate may be facilitating a complete rescue by a further mechanism(s) beyond anaplerosis. To examine this further, we performed an untargeted analysis using [U-^13^C]-pyruvate under GLS1 restriction and TGF-β_1_ stimulation and extended our labelling time to 48 hours to fully encompass all potential metabolites which may be mediating the rescue functions of pyruvate. This analysis identified 34 potential metabolites with high ^13^C enrichment and 27 of which were increased by pyruvate supplementation (Supplementary Data 2). Pathway analysis of these 27 metabolites revealed alanine, aspartate, and glutamate metabolism as the top pathway impact score, with glutamine catabolism and proline metabolism as second and third, respectively (Fig. 4a). Whilst we did detect malate and other TCA cycle intermediates in our untargeted analysis, the list of 27 metabolites was dominated by amino acids and acetylated amino acids. We therefore referenced our tracing data and analysed the peaks for glutamate, aspartate, proline and alanine. To allow for comparisons between the labels we first examined the isotopologue signal intensities to define total amounts contributed by either glucose or supplemented pyruvate following TGF-β_1_ stimulation and GLS1 inhibition. This analysis revealed that exogenous pyruvate increased (compared to Glc) the total amounts of glutamate, aspartate and proline derived from both PDH (m+2) and PC (m+3) under GLS1 inhibition and TGF-β_1_ stimulation (Fig. 4b-d). Pyruvate was also superior at generating alanine through GPT2 (indicative of the m+3 fraction) under these conditions than Glc-derived pyruvate (Fig. 4e). Examining total percent contributions of each label, we found that exogenous pyruvate sustained a greater fraction of the glutamate, aspartate and proline pools than Glc when GLS1 was inhibited (Fig. 4f-h). Alanine was unique in that over 70% was labelled from Glc by TGF-β_1_ and this was reduced to less than 5% by GLS1 inhibition (Fig. 4i). This strong decrease in carbon flux was likely due to both limiting availability of pyruvate (the carbon backbone of alanine) and glutamate (the amino group provider for alanine). GPT2 uses both of these substrates to generate alanine, and pyruvate flux through this enzyme was substantially increased by supplementation with pyruvate under CB-839 and TGF-β_1_ treated conditions, supporting roughly 20% of the total alanine pool (Fig. 4i). Out of these four amino acids, only alanine was significantly increased 48 hours into TGF-β_1_ signalling under GLS1 restriction with pyruvate present in the media (Fig. 4j and Supplementary Fig. 3e-g). Taken together, these data show that TGF-β_1_ stimulated cells have a greater ability to utilize exogenous pyruvate than glucose-derived pyruvate at increasing the isotopologue pool of glutamate, aspartate, proline and alanine following GLS1 inhibition.

**Figure 4.**
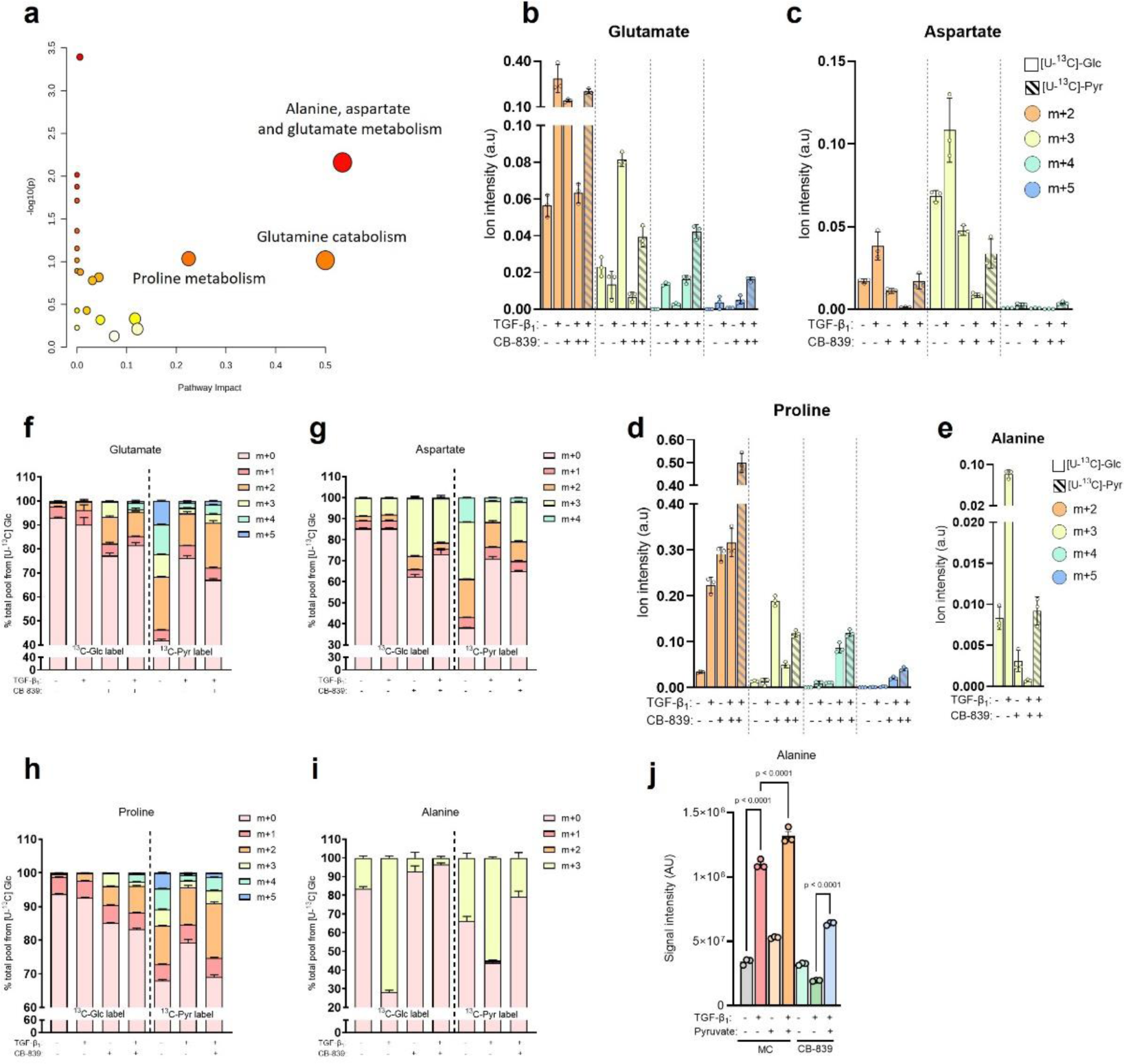
Exogenous pyruvate supports amino acid biosynthesis under GLS1 restriction. **a** Pathway analysis using MetaboAnalyst 5.0 of metabolites with high U-^13^C-pyruvate enrichment in 1 μM CB-839 treated and TGF-β_1_ (1 ng/ml) stimulated pHLFs following 48 h. **b-j** Intracellular isotopologue levels, **f-i** fractional enrichment and j total abundance of specified metabolite in pHLFs grown in DMEM^Low^supplemented with U-^13^C-glucose (5 mM) or U- ^13^C-pyruvate (1 mM) and pre-incubated with media control (0.1% DMSO) or 1 μM CB-839 for 1 h before stimulation with TGF-β_1_ (1 ng/ml) for 48 h and quantification achieved using LC-MS (*n*=3)

### 3.5 Pyruvate confers resistance to GLS1 inhibition through GDH-driven glutamate and GPT2-driven alanine biosynthesis

Our data so far strongly indicated that pyruvate was routing through the TCA cycle into glutamate to then bypass GLS1 inhibition. Our earlier results showed that both glutamate and α-ketoglutarate were capable of fully rescuing TGF-β_1_-induced collagen levels with CB-839 treatment (Fig. 2f). Given that α-ketoglutarate and glutamate are part of many bidirectional metabolic reactions, we questioned whether the cell-permeable form of α-ketoglutarate was mediating a rescue through regeneration of glutamate levels following GLS1 inhibition. Indeed, we found that the intracellular levels of not only glutamate but also those of alanine and proline were completely restored to TGF-β_1_ levels by α-ketoglutarate supplementation (Fig. 5a). We found that α-ketoglutarate supplementation did not regenerate aspartate levels which together with our silencing data of GOT1/GOT2 (Fig. 2c) suggested a dispensable role for aspartate biosynthesis in TGF-β_1_ induced collagen production.

**Figure 5.**
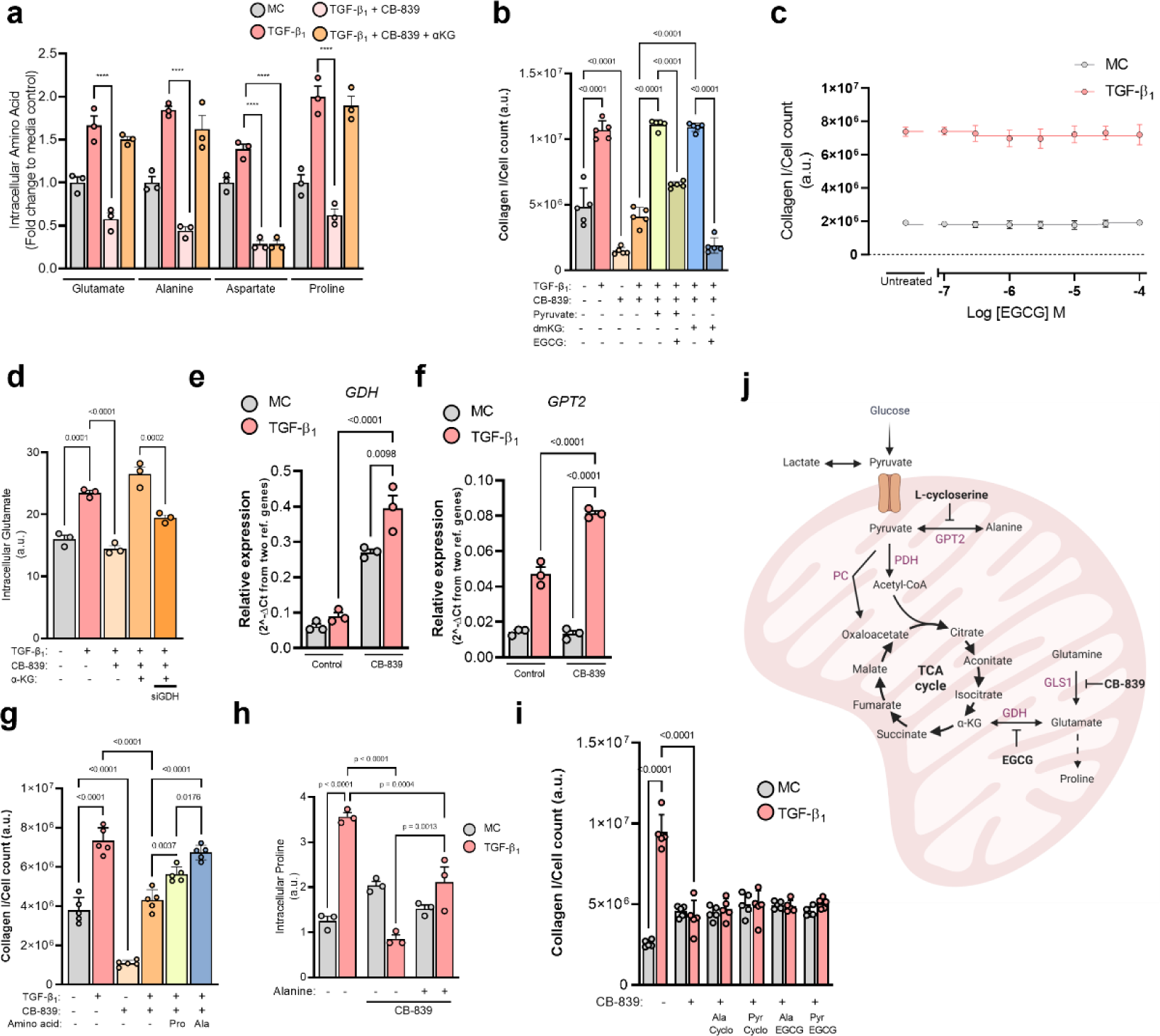
Pyruvate confers resistance to GLS1 inhibition through GDH-driven glutamate and GPT2-driven alanine biosynthesis. **a** Intracellular amino acid levels of pHLFs supplemented with dimethyl-a-ketoglutarate **(4 mM)** in the presence of 1 μM CB-839 and stimulated with TGF-β_1_ (1 ng/ml) for 48 h. b pHLFs were supplemented with dimethyl-a-ketoglutarate (4 mM) or pyruvate (1 mM) and pre-incubated with 1 μM CB-839 or 50 μM EGCG before stimulation with TGF-β_1_, (1 ng/ml) for 48 h and collagen I deposition assessed by macromolecular crowding assay and cell count using DAPI. c pHLFs were pre-incubated with increasing concentrations of EGCG for 1 h before being stimulated with TGF-β_1_ (1 ng/ml) for 48 h and collagen I deposition assessed by macromolecular crowding assay and normalised to cell count which was obtained by staining nuclei with DAPI. d pHLFs were transfected with non­targeting (NT) siRNA or siRNA targeting GDH, cells were pre-incubated with 1 μM CB-839 and dimethyl-a-ketoglutarate (4 mM) before stimulation with TGF-β_1_ (1 ng/ml) and intracellular glutamate levels quantified by HPLC after 48 hours. **e,f** pHLFs were treated with **1** μM CB-839 for 1 h prior to TGF-β_1_ (1 ng/ml) stimulation for 24 h and RNA quantified using qPCR **g** pHLFs were treated with 1 μM CB-839 for 1 **h** supplemented with alanine or proline (500 μM) prior to TGF-β_1_ (1 ng/ml) stimulation for 48 h and collagen I deposition assessed by macromolecular crowding assay and cell count using DAPI. h pHLFs were supplemented with alanine (500 μM) before stimulation with TGF-β_1_ (1 ng/ml) and CB-839 (1 μM)for48 h before being lysed and intracellular proline measured by HPLC (n=3). i pHLFs were treated as in **b** with alanine (500 pM) or pyruvate (1 mM) supplementation prior to a 1 h pre-incubation with 20 μM L-cycloserine or 50 μM EGCG before stimulation with TGF-β_1_ (1 ng/ml) for 48 h and collagen I deposition assessed by macromolecular crowding assay and cell count using DAPI. j Schematic showing pyruvate and glutamine routing into the TCA cycle. Data are presented as mean ± SD and differences evaluated between groups with two-way ANOVA with tukey multiple comparison testing.

Glutamate synthesis from α-ketoglutarate is largely mediated by aminotransferases, including GPT2 or GOT2, and glutamate dehydrogenase (GDH). The former requires the degradation of an amino acid and since GLS1 inhibition decreased amino acid levels this seemed unlikely to be the chosen pathway for glutamate synthesis following pyruvate supplementation. We therefore decided to inhibit GDH to determine if pyruvate and α-ketoglutarate were still capable of rescuing TGF-β_1_ induced collagen synthesis following CB-839 treatment. We found that inhibition of GDH, which is dispensable during TGF-β_1_-mediated collagen synthesis (Fig. 2c), prevented both pyruvate and α-ketoglutarate from rescuing collagen production under GLS1-restricted conditions (Fig. 5b,c). We observed that GDH inhibition decreased TGF-β_1_ collagen levels to a much lower extent with α- ketoglutarate than with pyruvate, suggesting that pyruvate primarily rescues CB-839 through amino acid synthesis rather than by fuelling the TCA cycle. We next measured intracellular glutamate levels under these conditions and found that silencing of GDH significantly decreased the regenerated levels of glutamate following α-ketoglutarate supplementation (Fig. 5d). These data showcase the resilience of the glutaminolytic axis and highlight that dual targeting of GLS1-derived glutamate and GDH-derived glutamate are required to effectively prevent TGF-β_1_-induced collagen synthesis. In further support of this, we found that inhibition of GLS1 led to an increase in both GPT2 and GDH mRNA levels, potentially indicating a physiologically relevant compensation mechanism (Fig. 5e,f)

We next individually introduced alanine and proline to examine whether CB-839-inhibited TGF-β_1_-induced collagen levels were restored. We found that proline induced a partial rescue whilst alanine was able to fully restore TGF-β_1_-induced collagen levels decreased by GLS1 inhibition (Fig. 5g). We therefore questioned how alanine might facilitate a complete rescue in GLS1-restricted conditions if the levels of proline, the biosynthetic pathway of which is crucial for TGF-β_1_ collagen production (Fig. 2c), were dependent on GLS1-derived glutamate (Supplementary Fig. 3g). We found that supplementation with alanine in CB-839 treated cells significantly increased TGF-β_1_ stimulated proline levels, indicating that alanine was capable of substituting for glutamate when GLS1 activity is inhibited (Fig. 5i). This would suggest a reversible function for GPT2 which degrades alanine to aminate α-ketoglutarate for glutamate generation. In support of this, we found that inhibiting GPT2 with L-cycloserine completely prevented alanine from rescuing GLS1 inhibition (Fig. 5h). Moreover, L-cycloserine also prevented pyruvate from rescuing GLS1 inhibition which highlights alanine biosynthesis as a major function of the pyruvate rescue (Fig. 5i,j).

### 3.5 Potential translational implications of targeting GLS1, GDH and GPT2 in lung fibrosis

From a therapeutic perspective, our data highlight two potential ways a GLS1 inhibitor strategy alone may be hampered in the setting of pulmonary fibrosis: (a) through pyruvate-sustained glutamate levels via GDH and (b) through upregulation of *GDH* and *GPT2* following GLS1 inhibition. With this in mind, we assessed the expression of the genes encoding these enzymes in the setting of IPF in publicly-available RNA-Seq datasets and found that all three genes are significantly elevated in IPF lung tissue compared with control lung (Fig. 6a-c) [27]. In order to determine if these genes are upregulated in pathological fibroblasts, we next analysed an IPF lung single-cell RNA-seq dataset and examined the significantly (p ≤ 0.05) upregulated genes in IPF myofibroblasts compared to control myofibroblasts [28]. This analysis identified 1,244 genes significantly increased in IPF myofibroblasts with subsequent pathway analysis revealing “glutamate and glutamine metabolism” as an enriched pathway (Fig. 6d). Other significant pathways identified were in line with expectations for a fibrotic disease and included for example “ECM-receptor interaction” and “protein processing”. Furthermore, “glutamate and glutamine metabolism” consisted of just two genes: *GLS1* and *GDH*, the expression of which were increased in IPF myofibroblasts. Interestingly, *GPT2* expression was low and did not pass significance cut-off. Furthermore, the glutamate-consuming and glutamine-synthesizing enzyme glutamine synthase (*GLUL*) was among the top downregulated genes in IPF myofibroblasts (60% reduction, p-value=7.2 x 10^-13^). This single-cell RNA-seq dataset was the first to identify four pathogenic fibroblasts subpopulations enriched in IPF, including a subpopulation of fibroblasts in the IPF lung characterized by high expression of hyaluronan synthase 1 *(HAS1*) and *COL1A1* expression (Fig. 6e). Expression values for both *GLS1* and *GDH* were highest in this subpopulation compared to the other three fibroblast groups identified in IPF (Fig. 6f,g).

**Figure 6.**
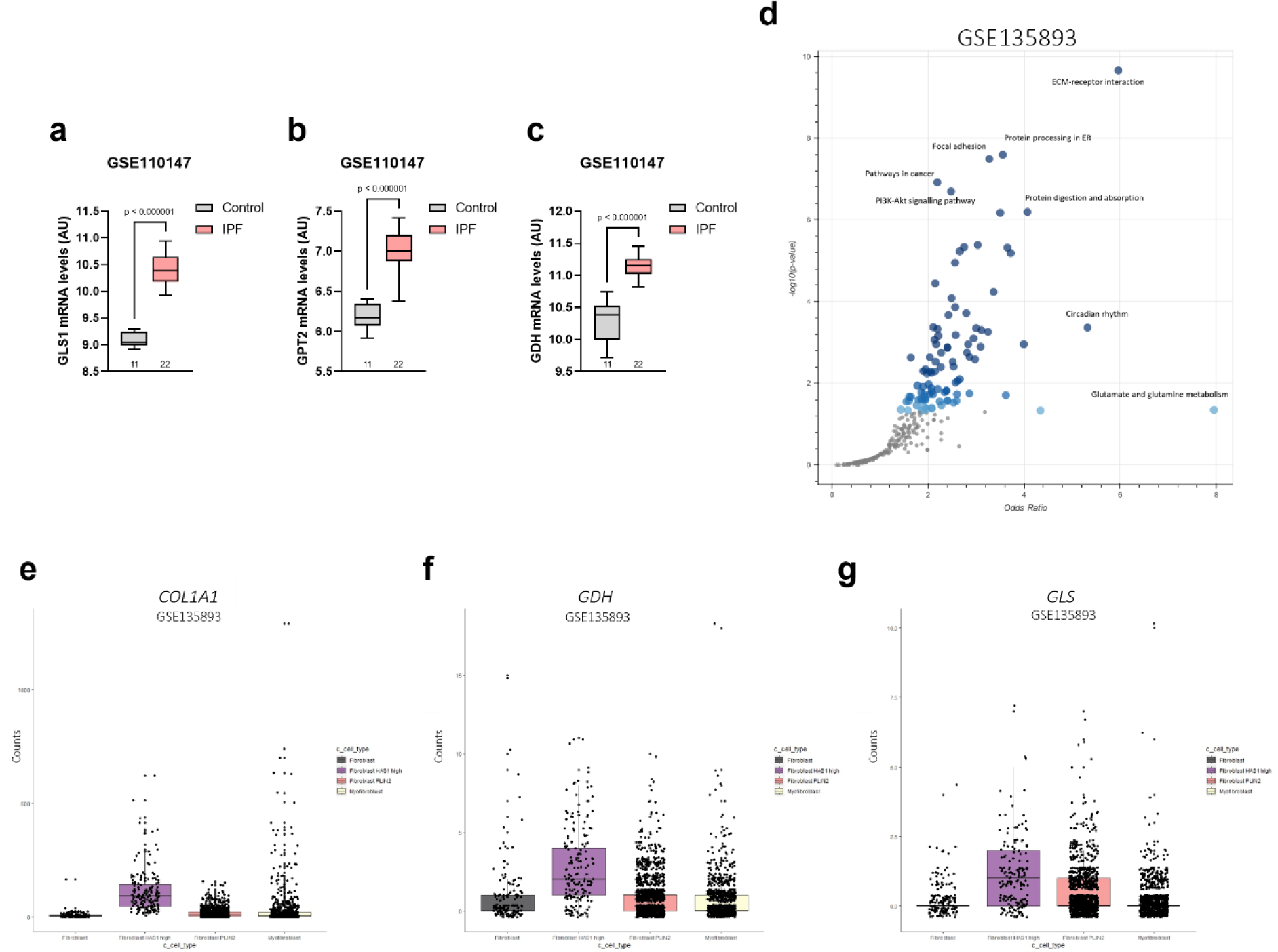
Potential translational implications of targeting GLS1, GDH and GPT2 in lung fibrosis. **a-c** mRNA expression of indicated genes in GSE110147 between a control lung and IPF lung. Patient numbers are indicated, **d** Pathway analysis (KEGG) of 1,244 upregulated genes (p < 0.05) derived from comparing control lung myofibroblasts to IPF lung myofibroblasts, **e-g** mRNA expression of indicated genes in GSE135893 comparing four fibroblast subpopulations found in IPF lung tissue.

Together, these data support the notion that the mechanistic metabolic axis investigated in the present report may contribute to the pathogenic production of collagen in the setting of fibrotic lung disease and further embolden a therapeutic strategy based on dual targeting of the glutamate-synthesizing enzymes GLS1 and GDH to exploit the metabolic vulnerabilities of hyper-secretory pathogenic fibroblast subpopulations (Fig. 7).

**Figure 7.**
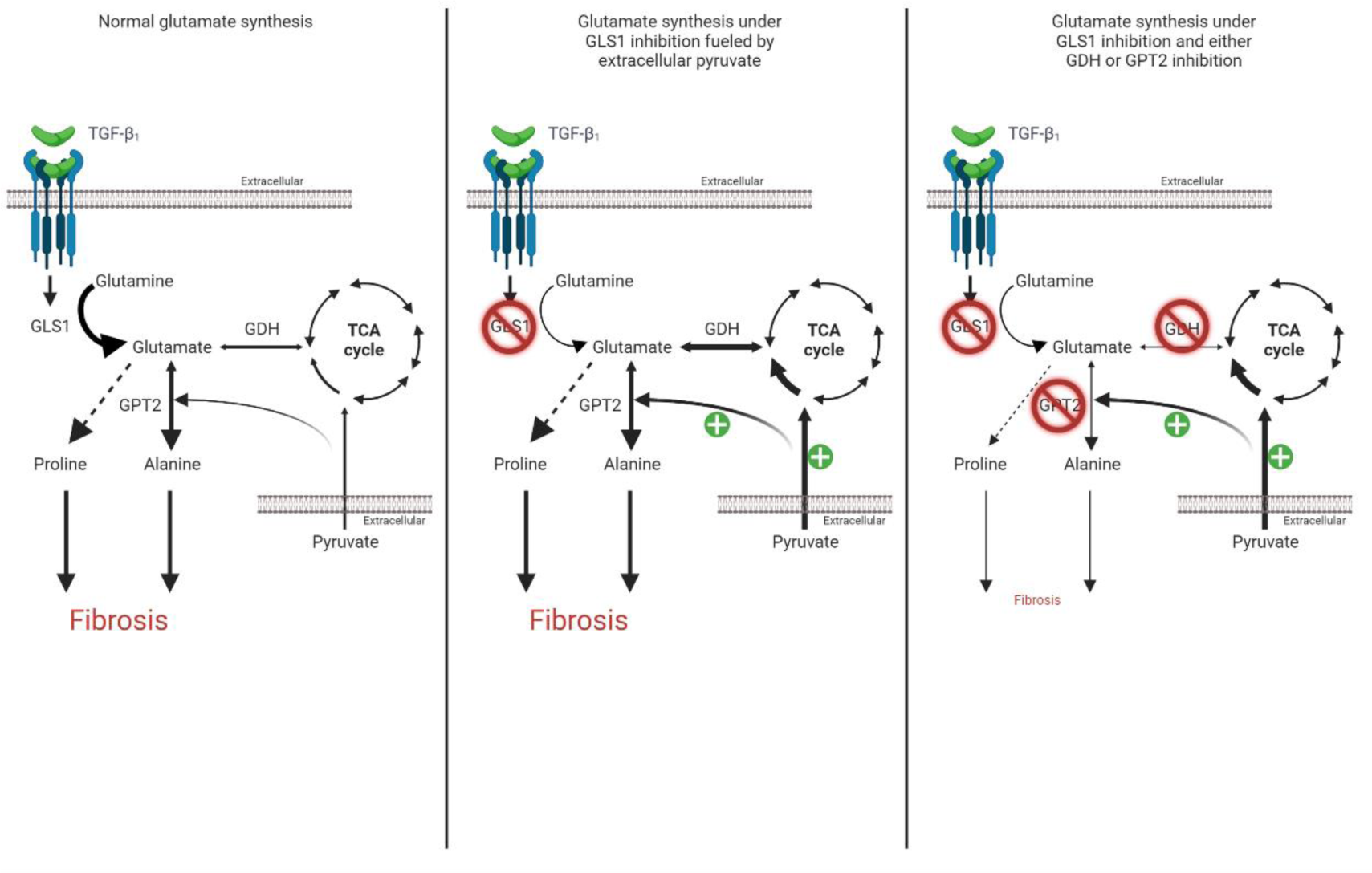
Proposed mechanism by which pyruvate routing under GLS1 inhibited conditions sustains fibrogenesis through GDH and GPT2. TGF-β_1_ increases GLS1 expression which leads to a conversion of intracellular glutamine levels to glutamate which supports alanine and proline biosynthesis, necessary for a TGF-β_1_ collagen response. Following GLS1 restriction, glutamate, alanine and proline levels decrease which are capable of being sustained by exogenous pyruvate which supplies glutamate through TCA cycle routing. Preventing this routing by inhibition of GDH or preventing alanine biosynthesis by inhibition of GPT2 are two strategies to effectively limit the TGF-β_1_ collagen response under GLS1 restriction. (TGF-β_1_: Transforming growth factor p,; GLS1: Glutaminase 1; GDH: Glutamate dehydrogenase; GPT2: Glutamic-pyruvic transaminase 2).

## 4 Discussion

Fibroblasts play a central role in the turnover of the ECM in homeostasis and during tissue repair. In fibrosis, this turnover becomes dysregulated and culminates in the progressive accumulation of ECM proteins, primarily collagens, which leads to the obliteration of tissue architecture and eventual organ failure. This pathological process has also been implicated in promoting tumour growth as part of the stromal reaction in cancer [17]. In this study, we report that media composition, particularly changes in the extracellular concentrations of glucose, glutamine and pyruvate, have a major influence on the global TGF-β_1_ regulated transcriptome and intracellular amino acid profile in primary human lung fibroblasts. We further uncover a critical role for pyruvate metabolism in influencing the impact of glutaminolysis inhibition on the TGF-β_1_-induced collagen response. We trace this to a GDH-dependent glutamate and a GPT2-dependent alanine biosynthetic axis. In addition to furthering our fundamental understanding of glycolysis during the TGF-β_1_-induced fibrogenic response, these findings have important implications for the development of novel therapeutic approaches aimed at targeting the metabolic vulnerabilities of hyper-synthetic fibroblasts as a novel anti-fibrotic strategy.

Media composition greatly impacted the global transcriptional response to TGF-β_1_, both in terms of the number of genes impacted as well as the magnitude of the response. Whilst we did not individually examine the influence of glucose, glutamine and pyruvate concentration differences on the TGF-β_1_ transcriptome, there is evidence that high carbon load (glucose and pyruvate) can affect the cellular epigenome through alteration of chromatin acetylation and methylation sites, which in turn facilitates transcription factor accessibility to a greater number of genes [29, 30]. The major acetylation substrate, acetyl-CoA has recently been described as a key metabolic regulator of the epigenome and our untargeted data presented here identified a large number of acetyl-CoA conjugated metabolites enriched in the presence of pyruvate. Furthermore, our labelling study showed that pyruvate supplementation greatly increased TGF-β_1_-stimulated acetyl-CoA levels. It is therefore tempting to speculating that the presence of extracellular pyruvate may influence transcriptional responses via an epigenetic modifying mechanism.

In this study we focussed on exploring the contribution of media composition in terms of the anti-fibrotic potential of CB-839, a GLS1 inhibitor currently under investigation in several cancer trials. This compound was found to be effective in halting the TGF-β_1_ collagen response in conditions of limited pyruvate or alanine availability and was partially effective in conditions of proline availability or insufficient glycolytic flux leading to pyruvate utilisation in the mitochondria. We traced the effects of extracellular pyruvate in mediating resistance to GLS1 inhibition during TGF-β_1_-induced collagen synthesis to its ability to act as a substrate for alanine biosynthesis and, to a lesser extent, proline biosynthesis. Pyruvate concentration is often undefined in conventional cell culture media and glucose and glutamine are often set at supraphysiological concentrations-making it difficult to compare findings across different studies. We found that supraphysiological levels of glutamine, glucose and pyruvate led to a higher pool of intracellular alanine which may explain why other studies using conventional fibroblast media may have underestimated the importance of GPT2 as a critical enzyme for TGF-β_1_-driven collagen production [31, 32]. In the cancer setting, GPT2 and GLS1 were identified following a CRISPR screen as critical enzymes for cell viability dependent on extracellular alanine and pyruvate availability, respectively [16]. To further confound cross-analysis of relevant fibrosis studies, the quantification of collagen can be performed using various technologies which measure collagen synthesis at different steps during the biosynthetic process. For example, CB-839 has been reported to show some efficacy in inhibiting intracellular collagen I levels following TGF-β_1_ stimulation *in vitro* and partially inhibit total lung collagen levels in experimental lung fibrosis models [24]. Our data shows that the anti-fibrotic effect of GLS-1 inhibition is highly dependent on the availability of key carbon and nitrogen sources and the inhibitory effect is enhanced with additional targeting of pyruvate-dependent alanine (GPT2) or glutamate synthesis enzymes (GDH).

The link between pyruvate and glutamine metabolic pathways has garnered much interest in the oncology setting, where pyruvate has been shown to sustain the CB-839-depleted TCA cycle in proliferating cancer cells [33]. Whilst we did find that the levels of TCA cycle intermediates following TGF-β_1_ stimulation were dependent on GLS1 in fibroblasts, we found a limited impact on these levels following pyruvate supplementation and found that the pyruvate-mediated rescue of CB-839 treatment on enhanced collagen levels was dependent on GDH-driven glutamate generation, a reaction stemming from α-ketoglutarate. This notion is further supported by a recent study examining GLS1 inhibition in lung fibroblasts which showed no reduction in oxygen consumption (indicative of electron transport chain (ETC) activity and, by proxy, of TCA cycle activity) following CB-839 treatment [31]. Furthermore, we have previously published on the dispensable nature of the ETC for lung fibroblasts to mount a full TGF-β_1_ induced collagen response [34]. Taken together the evidence might now be mounting that biosynthetic processes might be more limiting than bioenergetic processes (such as NADH generation through the TCA cycle) for TGF-β_1_-enhanced collagen production.

Our results highlight alanine as a critical amino acid supporting TGF-β_1_-induced collagen synthesis. In the absence of extracellular alanine, siRNA-mediated silencing or pharmacological inhibition of GPT2 completely prevented pHLFs from mounting a full collagen response following TGF-β_1_ exposure. Of note, GPT2 knockdown did not affect baseline collagen levels which suggests that targeting GPT2 therapeutically would preferentially limit collagen production in TGF-β_1_-activated fibroblasts. Alanine derived from the circulation may be a factor to consider for such a therapeutic approach and could be modulated to limit fibrogenesis, as has been explored in the setting of cancer cell proliferation by intravenous administration of amino acid degrading enzymes, such as asparaginase or cysteinase [35, 36].

Taken together, the findings presented herein shed important light on the complex interplay between metabolic networks and highlight a therapeutic strategy aimed at targeting both mitochondrial pyruvate metabolism and glutaminolysis to interfere with the pathological deposition of collagen in the setting of pulmonary fibrosis and potentially other fibrotic conditions.

## Abbreviations

ALDH18A1: aldehyde dehydrogenase 18 family member A1
ASNS: asparagine synthetase
DCA: dichloroacetate
ECM: extracellular matrix
EGCG: epigallocatechin gallate
ETC: electron transport chain
GDH: glutamate dehydrogenase
GLS1: glutaminase 1
GOT1: glutamic-oxaloacetic transaminase 1
GOT2: glutamic-oxaloacetic transaminase 2
GPT2: glutamic-pyruvic transaminase 2
GSH: glutathione
P5CS: pyrroline-5-carboxylate synthase
PC: pyruvate carboxylase
PDH: pyruvate dehydrogenase
pHLFs: primary human lung fibroblasts
PSAT1: phosphoserine aminotransferase 1
SHMT2: serine hydroxymethyltransferase2
TCA: tricarboxylic acid cycle
TGF-β_1_: transforming growth factor β_1_

## Acknowledgements

The authors gratefully acknowledge Dr Silvia Parolo from the Centre for Computational and Systems Biology (COSBI) for her contributions to the analysis of the single-cell RNAseq datset GSE135893.

## Author contributions

GC conceptualised the approach, performed experiments, interpreted the data, generated the figures and collaboratively drafted the manuscript. RCC conceived the study, secured the funding and supervised the work. JAMW and GC performed the bioinformatics analysis of RNA-Seq datasets. MP and DG performed experiments and provided RNA-Seq datasets. BS helped interpret data and edited the manuscript. VM performed the LC-MS analysis for the metabolomics study under the supervision of KB who also interpreted data. VP and MP provided expert input into the study. All authors reviewed and approved the final submitted manuscript.

## Supplementary figures

**Supplementary Figure 1.**
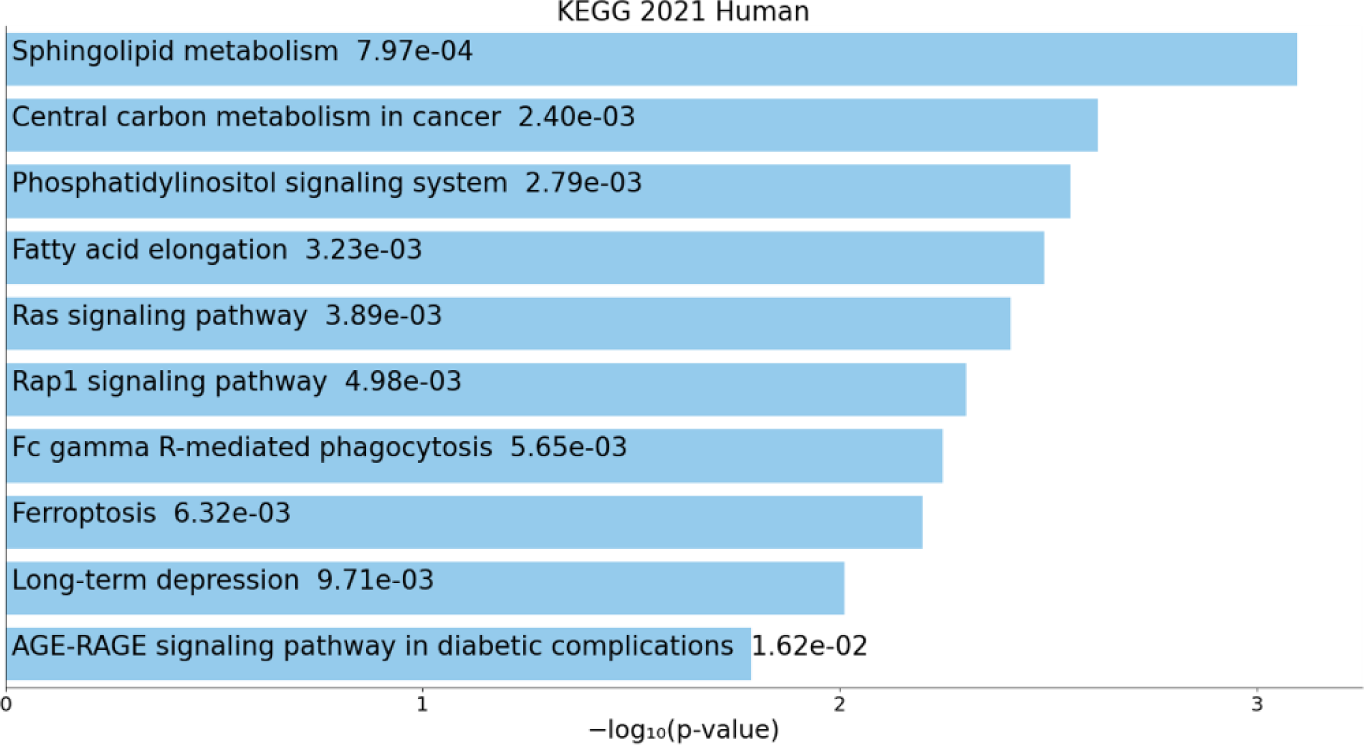
KEGG pathway analysis of DEGs unique to pHLFs cultured in DMEM^High^. DEGs derived from RNA-Seq analysis of pHLFs 24 hours following TGF-β_1_ (1 ng/ml) stimulation in DMEM^High^ and DEGs found in DMEM^Low^ were removed, leaving DEGs unique to DMEM^High^.

**Supplementary Figure 2.**
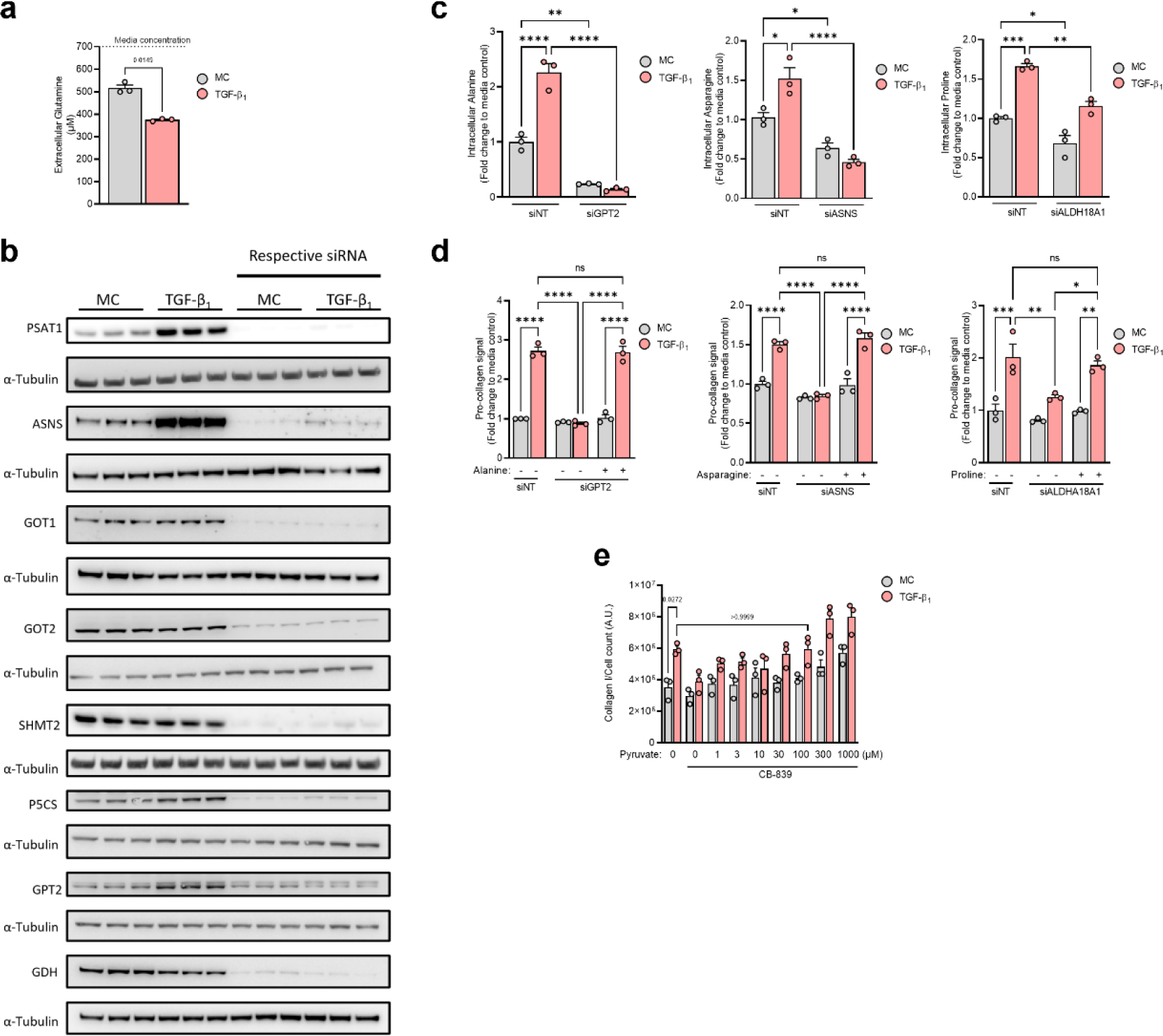
Metabolite rescues of TGF-β_1_ induced collagen under metabolic enzyme inhibition or protein expression knockdown. **a** Extracellular glutamine levels 48 hours following TGF-β_1_ (1 ng/ml) stimulation in DMEM^Low^. **b** pHLFs were transfected with non-targeting (NT) siRNA or siRNA targeting PSAT1, ASNS, GOT1, GOT2, SHMT2, ALDH18A1 (P5CS), GPT2 or GDH and protein expression measured by immunoblot 24 hours following TGF-β_1_ (1 ng/ml). **c, d** pHLFs were stimulated with TGF-β_1_ (1 ng/ml) for 48 h following transfection with non-targeting siRNA for control or siRNA targeting GPT2, ASNS or ALDH18A1 and intracellular levels of alanine, asparagine or proline (respectively) quantified using HPLC. **d** Media supplemented with respective metabolite (500 pM) and hydroxyproline quantified using HPLC (*n*=3) **e** pHLFs were grown in DMEM^Low^ and pre-incubated with increasing concentrations of pyruvate for 1 h before being stimulated with TGF-β_1_ (1 ng/ml) and treated with CB-839 (1 pM) for 48 h and collagen I deposition assessed by macromolecular crowding assay. Data are expressed as collagen I signal as a fold-change of the media control (0.1% DMSO) normalised to cell count which was obtained by staining nuclei with DAPI. Data are presented as mean ± SD and differences evaluated between groups with two-way ANOVA with tukey multiple comparison testing.

**Supplementary Figure 3.**
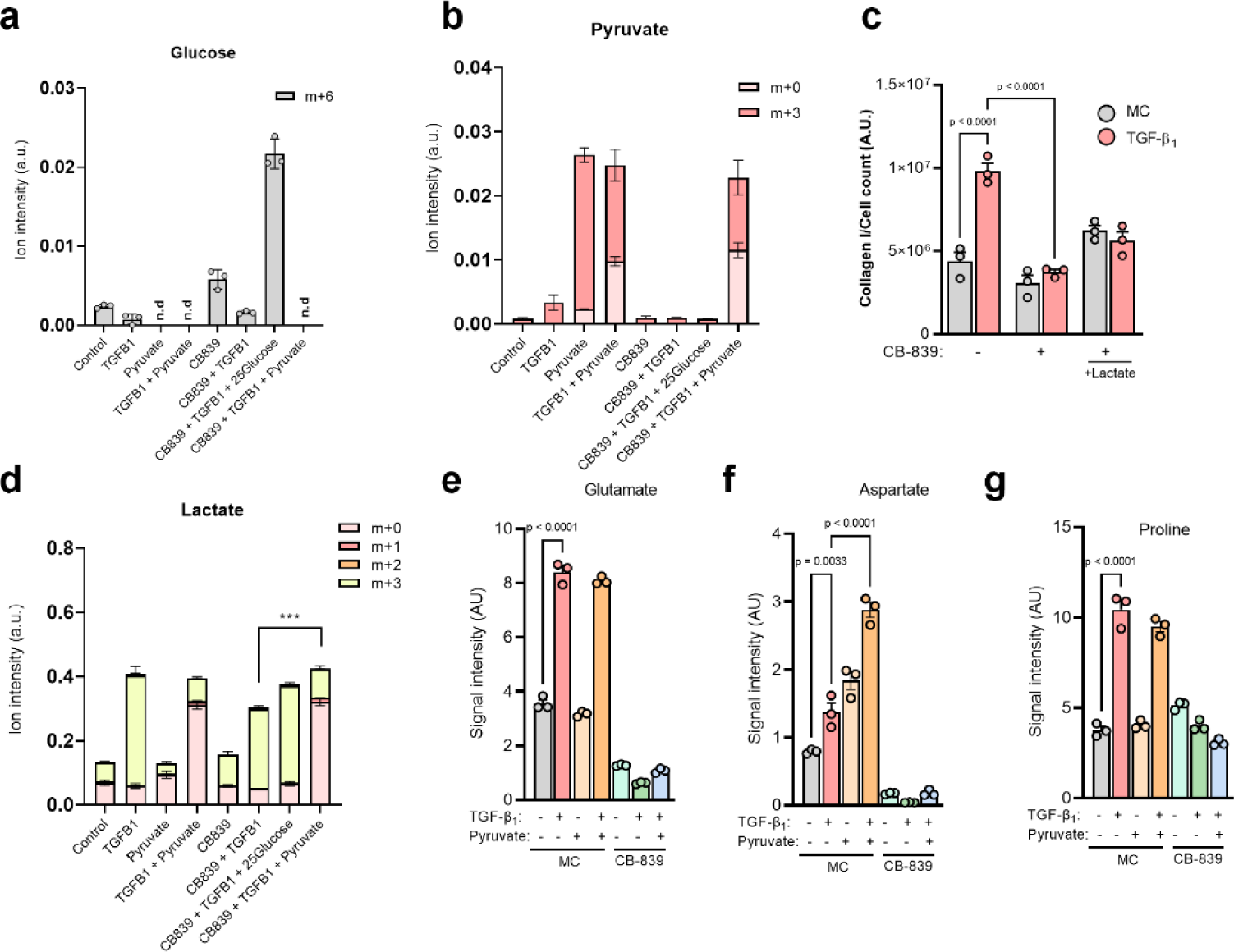
High glucose does not phenocopy exogenous pyruvate. **a,b and d-g** Intracellular isotopologue levels and **e-g** total abundance of specified metabolite in pHLFs grown in DMEM^Low^ supplemented with U-^13^C-glucose (5 mM) or U-^13^C-pyruvate (1 mM) and pre-incubated with media control (0.1% DMSO) or 1 pM CB-839 for 1 h before stimulation with TGF-β_1_ (1 ng/ml) for 48 h and quantification achieved using LC-MS (*n*=3). c Collagen deposition quantified 48 hours after TGF-β_1_ (1 ng/ml) stimulation from pHLFs growing in DMEM^Low^ and 1 pM CB-839 with supplementation of lactate (10 mM). Data are presented as mean ± SD and differences evaluated between groups with two-way ANOVA with tukey multiple comparison testing.

