## Supplementary figures for "Pyruvate metabolism dictates fibroblast sensitivity to GLS1 inhibition during fibrogenesis"

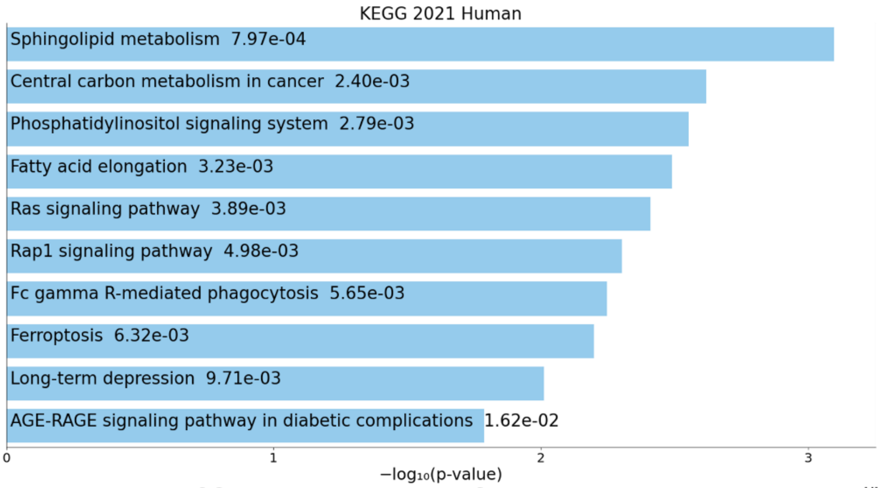


**Supplementary Figure 1. KEGG pathway analysis of DEGs unique to pHLFs cultured in DMEM$^{High}$.** DEGs derived from RNA-Seq analysis of pHLFs 24 hours following TGF-$\beta_1$ (1 ng/ml) stimulation in DMEM$^{High}$ and DEGs found in DMEM$^{Low}$ were removed, leaving DEGs unique to DMEM$^{High}$.

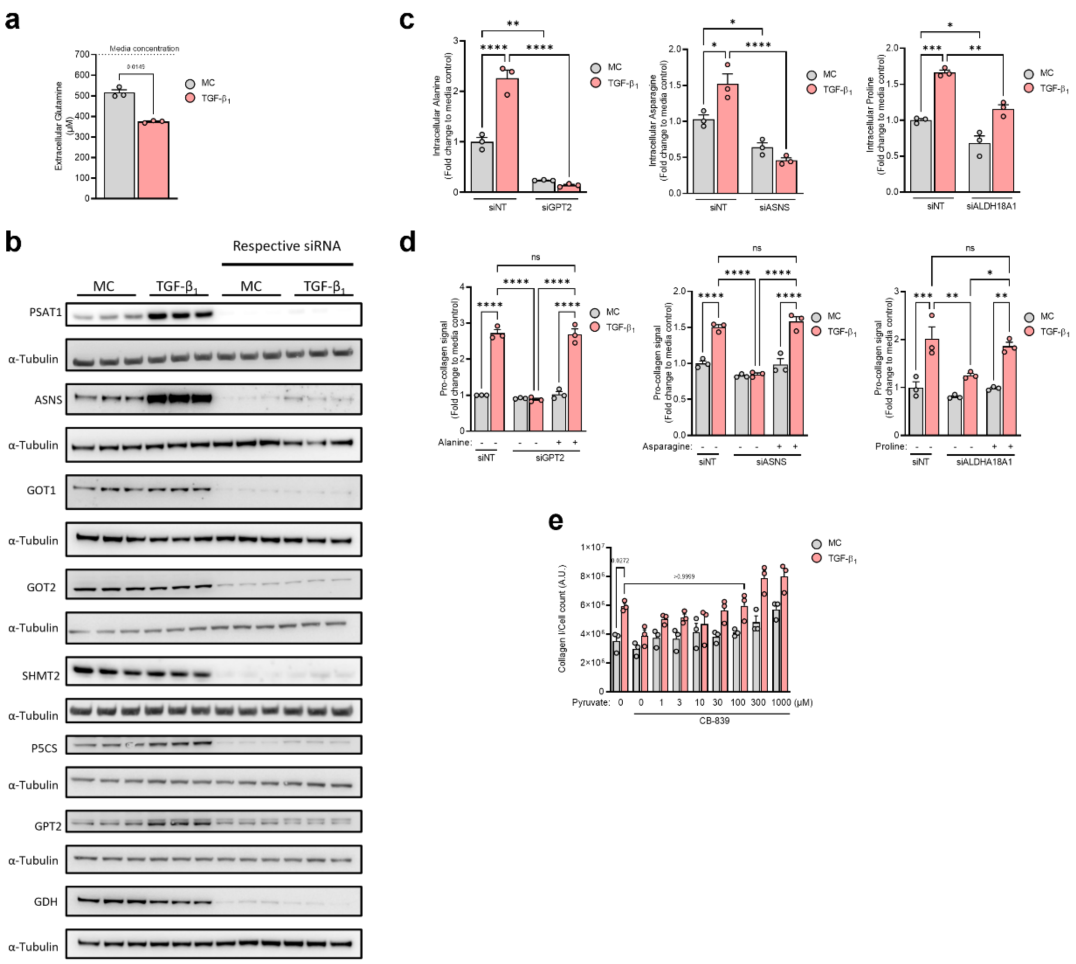


**Supplementary Figure 2. Metabolite rescues of TGF-$\beta_1$-induced collagen under metabolic enzyme inhibition or protein expression knockdown.** **a** Extracellular glutamine levels 48 hours following TGF-$\beta_1$ (1 ng/ml) stimulation in DMEM$^{Low}$. **b** pHLFs were transfected with non-targeting (NT) siRNA or siRNA targeting PSAT1, ASNS, GOT1, GOT2, SHMT2, ALDH18A1 (P5CS), GPT2 or GDH and protein expression measured by immunoblot 24 hours following TGF-$\beta_1$ (1 ng/ml). **c, d** pHLFs were stimulated with TGF-$\beta_1$(1 ng/ml) for 48 h following transfection with non-targeting siRNA for control or siRNA targeting GPT2, ASNS or ALDH18A1 and intracellular levels of alanine, asparagine or proline (respectively) quantified using HPLC. **d** Media supplemented with respective metabolite (500 μM) and hydroxyproline quantified using HPLC (*n*=3). **e** pHLFs were grown in DMEM$^{Low}$ and pre-incubated with increasing concentrations of pyruvate for 1 h before being stimulated with TGF-$\beta_1$ (1 ng/ml) and treated with CB-839 (1 μM) for 48 h and collagen I deposition assessed by macromolecular crowding assay. Data are expressed as collagen I signal as a fold-change of the media control (0.1% DMSO) normalised to cell count which was obtained by staining nuclei with DAPI. Data are presented as mean ± SD and differences evaluated between groups with two-way ANOVA with tukey multiple comparison testing.

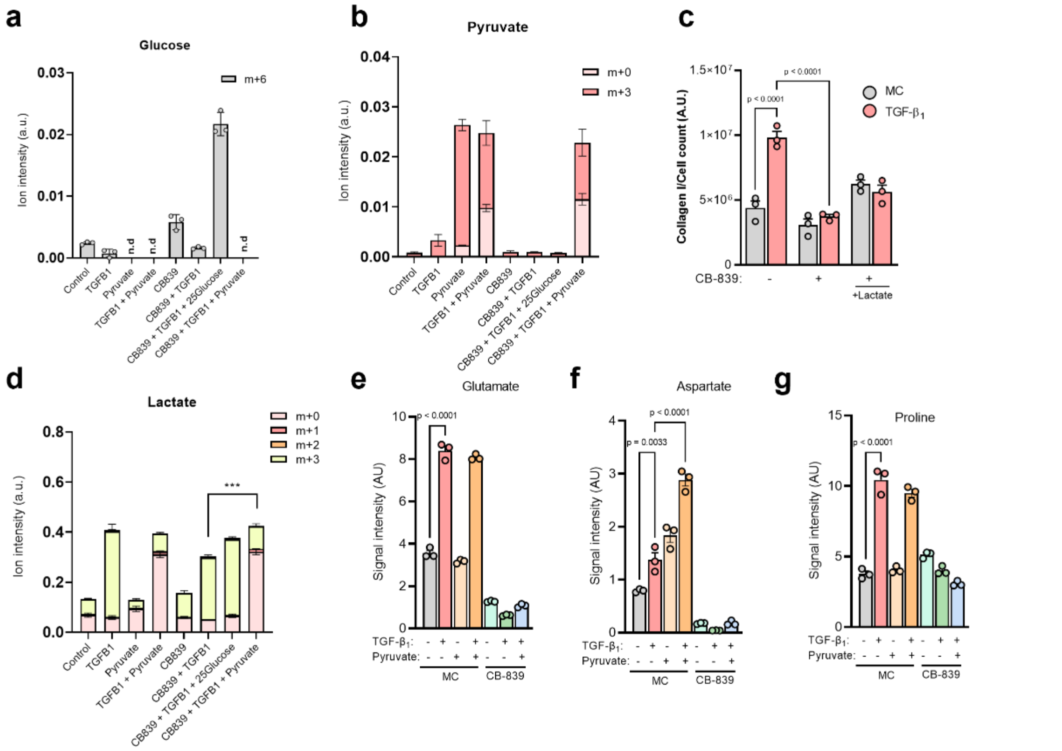


**Supplementary Figure 3. High glucose does not phenocopy exogenous pyruvate. a,b and d-g** Intracellular isotopologue levels and **e-g** total abundance of specified metabolite in pHLFs grown in DMEM$^{Low}$ supplemented with U-$^{13}$C-glucose (5 mM) or U-$^{13}$C-pyruvate (1 mM) and pre-incubated with media control (0.1% DMSO) or 1 μM CB-839 for 1 h before stimulation with TGF-$\beta_1$ (1 ng/ml) for 48 h and quantification achieved using LC-MS (*n*=3). c Collagen deposition quantified 48 hours after TGF-$\beta_1$ (1 ng/ml) stimulation from pHLFs growing in DMEM$^{Low}$ and 1 μM CB-839 with supplementation of lactate (10 mM). Data are presented as mean ± SD and differences evaluated between groups with two-way ANOVA with tukey multiple comparison testing.
